# Evolved sequence features within the intrinsically disordered tail influence FtsZ assembly and bacterial cell division

**DOI:** 10.1101/301622

**Authors:** Megan C. Cohan, Ammon E. Posey, Steven J. Grigsby, Anuradha Mittal, Alex S. Holehouse, Paul J. Buske, Petra A. Levin, Rohit V. Pappu

**Affiliations:** Department of Biomedical Engineering and Center for Biological Systems Engineering, St. Louis, St. Louis, MO 63130; Department of Biology, Washington University in St. Louis, St. Louis, MO 63130

## Abstract

Intrinsically disordered regions (IDRs) challenge the well-established sequence-structure-function paradigm for describing protein function and evolution. Here, we direct a combination of biophysical and cellular studies to further our understanding of how the intrinsically disordered C-terminal tail of FtsZ contributes to cell division in rod-shaped bacteria. FtsZ is a modular protein that encompasses a conserved GTPase domain and a highly variable intrinsically disordered C-terminal tail (CTT). The CTT is essential for forming the cytokinetic Z-ring. Despite poor sequence conservation of the CTT, the patterning of oppositely charged residues, which refers to the extent of linear mixing / segregation of oppositely charged residues within CTT sequences is bounded within a narrow range. To assess the impact of evolutionary bounds on charge patterning within CTT sequences we performed experiments, aided by sequence design, to quantify the impact of changing the patterning of oppositely charged residues within the CTT on the functions of FtsZ from *B*. *subtilis*. Z-ring formation is robust if and only if the extent of linear mixing / segregation of oppositely charged residues within the CTT sequences is within evolutionarily observed bounds. Otherwise, aberrant, CTT-mediated, FtsZ assemblies impair Z-ring formation. The complexities of CTT sequences also have to be above a threshold value because FtsZ variants with low complexity CTTs are not tolerated in cells. Taken together, our results suggest that CTT sequences have evolved to be “just right” and that this is achieved through an optimal extent of charge patterning while maintaining the sequence complexity above a threshold value.

## Introduction

Intrinsically disordered regions (IDRs) feature prominently in eukaryotic proteins (Dunker et al., 2002a; van der Lee et al., 2014). These regions are associated with a range of molecular functions that contribute to an assortment of cellular processes (Dunker et al., 2002b; Iakoucheva et al., 2002; Iakoucheva et al., 2004; Wright and Dyson, 2015). IDRs and intrinsically disordered proteins (IDPs) defy the well-established sequence-structure-function paradigm that provides a coherent framework for connecting sequence-encoded information to molecular functions for proteins that fold into well-defined three-dimensional structures (Babu, 2016). This is because conformational heterogeneity is a defining hallmark of IDRs / IDPs (Babu et al., 2012; Forman-Kay and Mittag, 2013; Lyle et al., 2013; Wright and Dyson, 2015). Despite this sequence-intrinsic conformational heterogeneity it has become clear that IDRs and IDPs have definable sequence-to-conformation relationships (Das et al., 2015; Forman-Kay and Mittag, 2013; Gibbs and Showalter, 2016). The overall sizes, shapes, and amplitudes of conformational fluctuations of IDRs / IDPs are governed by compositional parameters such as the fraction of charged residues and the overall proline contents (Hofmann et al., 2012; Holehouse et al., 2017; Mao et al., 2013; Muller-Spath et al., 2010). Additional contributions to sequence-to-conformation relationships include the linear patterning of oppositely charged residues as well as the linear patterning of proline and charged residues (Das and Pappu, 2013; Martin et al., 2016).

Progresses in advancing our understanding of sequence-to-conformation relationships of IDRs / IDPs (Mollica et al., 2016; Schneider et al., 2015) have opened the door to using sequence design as a tool to investigate the functional and phenotypic effects of modulating sequence-to-conformation relationships of IDPs / IDRs (Das et al., 2016; Sherry et al., 2017; Staller et al., 2018). These studies have been directed at yeast proteins that control cell cycle arrest (Das et al., 2016), mating type switching (Martin et al., 2016), and transcriptional activation (Staller et al., 2018) and mammalian systems that regulate cell differentiation (Sherry et al., 2017), tumor suppression (Borcherds et al., 2014), and transcriptional regulation (Portz et al., 2017). There is also growing interest in the roles of IDRs / IDPs as modulators of the driving forces for the formation of membraneless organelles and protein-RNA bodies in eukaryotic systems (Dao et al., 2018; Harmon et al., 2017; Hernandez-Vega et al., 2017; Lin et al., 2017; Lin et al., 2018; Mitrea et al., 2018; Mitrea and Kriwacki, 2016; Wei et al., 2017).

In contrast to eukaryotic systems and viruses, where the functional importance of IDRs / IDPs is reasonably well established (Berlow et al., 2017; Jensen et al., 2011; Mollica et al., 2016; Wright and Dyson, 2015), the prevailing dogma is that bacterial proteins largely conform to the classical sequence-structure-function paradigm (Dunker et al., 2015; Pavlovic-Lazetic et al., 2011). This view has emerged from bioinformatics analysis, which shows that only a small percentage of bacterial proteins include long IDRs (Brown et al., 2011; Chen et al., 2006; Yruela et al., 2017). The apparent bias against IDRs / IDPs in bacterial proteomes has been ascribed to a variety of factors. These include shorter protein lengths, the importance of well-defined structure in metabolic enzymes, and the purportedly focused / limited repertoire of functions associated with bacterial proteins (Brown et al., 2010; Dunker et al., 2001; Yruela et al., 2017). However, recent studies have demonstrated that while IDRs / IDPs may make up a small fraction of proteins / protein regions in bacterial proteomes when compared to eukaryotic ones, the synergies between IDRs and folded domains contribute directly to an assortment of functions. Prominent bacterial IDRs / IDPs include proteins involved in regulating cell division (Buske and Levin, 2013; Buske et al., 2015), DNA replication (Kozlov et al., 2015; Shereda et al., 2008), protein and RNA quality control (Ait-Bara and Carpousis, 2015; Ait-Bara et al., 2015; Foit et al., 2013; Jakob et al., 2014; Quan et al., 2014; Reichmann et al., 2012; Updegrove et al., 2016), bacterial warfare (Bonsor et al., 2008; Hecht et al., 2009; Papadakos et al., 2015), biofilm formation (Gruszka et al., 2016; Gu et al., 2017; Whelan and Potts, 2015), and chemotaxis (Daughdrill et al., 1997; Dedmon et al., 2002; Ding et al., 2009). These findings raise a key question: Do the specific sequence-to-conformation relationships uncovered for archetypal eukaryotic IDRs / IDPs have any bearing on the functions and phenotypes that are influenced by IDRs / IDPs in bacteria? To answer this question, we have focused our current study on the C-terminal disordered tail of FtsZ, a protein that is directly involved in regulating cell division in rod-shaped bacteria (Buske and Levin, 2013; Buske et al., 2015).

Cell division across all domains of life is initiated by the formation of a cytokinetic ring of key cytoskeletal proteins at the nascent division site (Addi et al., 2018; Baluska et al., 2006; Guizetti and Gerlich, 2010; Huckaba and Pon, 2002; Margolin, 2001; Rincon and Paoletti, 2016; Straight and Field, 2000; Thieleke-Matos et al., 2017; Tolliday et al., 2001). The cytokinetic ring serves as the foundation for assembly of the cell division machinery. In animals and fungi the cytokinetic ring is composed of the ATPases actin and myosin (Addi et al., 2018). In rod-shaped bacteria the main building block of the cytokinetic ring, also known as the Z-ring, is the essential GTPase FtsZ, which is a homolog of eukaryotic tubulin (Buske et al., 2015; Erickson et al., 2010; Huang et al., 2013; Janakiraman and Goldberg, 2004; Lutkenhaus, 1993; Meier and Goley, 2014; Rothfield et al., 1999).

While monomeric FtsZ can bind GTP, the active site for GTP hydrolysis is formed at the interface between two FtsZ subunits (Oliva et al., 2007). FtsZ forms single-stranded polymers *in vitro* and GTP-binding promotes FtsZ polymerization (Buske et al., 2015). FtsZ polymers serve as a treadmilling platform for the division machinery, particularly enzymes required for synthesis of peptidoglycans (Bisson-Filho et al., 2017; Monteiro et al., 2018). In cells, the FtsZ ring is highly dynamic with subunit turnover occurring on the order of seconds (Anderson et al., 2004). *In vitro* experiments indicate that FtsZ polymerization and Z-ring formation involves distinct transitions (Bi and Lutkenhaus, 1991; Erickson et al., 2010). In the presence of millimolar concentrations of GTP, FtsZ forms single-stranded polymers or filaments (Mukherjee and Lutkenhaus, 1994). This type of polymerization can be described as an isodesmic process (Skillman et al., 2013), which is characterized by similar association constants for the initial dimerization step and the elongation of polymers (Caplan and Erickson, 2003; Romberg et al., 2001). Filaments can associate laterally with one another to form bundles (Gonzalez et al., 2005; Romberg and Levin, 2003). GTP hydrolysis promotes the bending of laterally associated polymers to form wreath-like structures that vary in thickness around the circumference (den Blaauwen et al., 2017).

Calorimetric measurements suggest that the formation of FtsZ assemblies requires the crossing of a critical concentration threshold (Caplan and Erickson, 2003; Chen et al., 2005). This apparent cooperativity is at odds with the observation that FtsZ forms stable singlestranded filaments, which is the defining characteristic of isodesmic assembly (Oosawa and Kasai, 1962). Numerous models have been developed to describe how FtsZ assembles cooperatively. These models suggest that FtsZ subunits might undergo conformational changes to facilitate the population of a high affinity state that favors polymerization (Miraldi et al., 2008). Detailed molecular and structural evidence for such a high affinity state remains elusive. In this context, it is worth noting that most models for FtsZ assembly have been motivated primarily by the structure and dynamics of the conserved and well-folded GTPase domain.

We propose that clues to the observed complexities and regulation of FtsZ assembly come from the distinctive protein architecture, which include modules other than the GTPase domain (Figure 1A). The modular architecture of FtsZ encompasses a folded N-terminal core domain that forms an active GTP hydrolysis site upon dimerization. The negatively charged core includes the GTP binding site and the coordinating T7 loop. The interface between the nucleotide binding site and the T7 loop (in dimers and higher order polymers) forms the active site for GTP hydrolysis (Oliva et al., 2007). The C-terminal stretch of FtsZ includes an intrinsically disordered linker (CTL), a 17-residue C-terminal peptide (CTP), and 7-10 residue stretch that is rich in basic residues and has been designated as the C-terminal variable region or CTV (Buske and Levin, 2012, 2013; Buske et al., 2015). The CTP is an alpha-helical molecular recognition element (Oldfield et al., 2005) that coordinates heterotypic protein-protein interactions involving FtsZ. The interaction partners that engage with FtsZ through the CTP include the protein FtsA, which assists in the bundling of FtsZ protofilaments and anchoring of Z-rings to bacterial membranes, and SepF, which is required for the late stages of cell division (Adams and Errington, 2009). Depending on its charge, the CTV can play a significant role in mediating lateral interactions between FtsZ polymers *in vitro* and in determining the integrity of the FtsZ ring *in vivo* (Buske and Levin, 2012).

**Figure 1:**
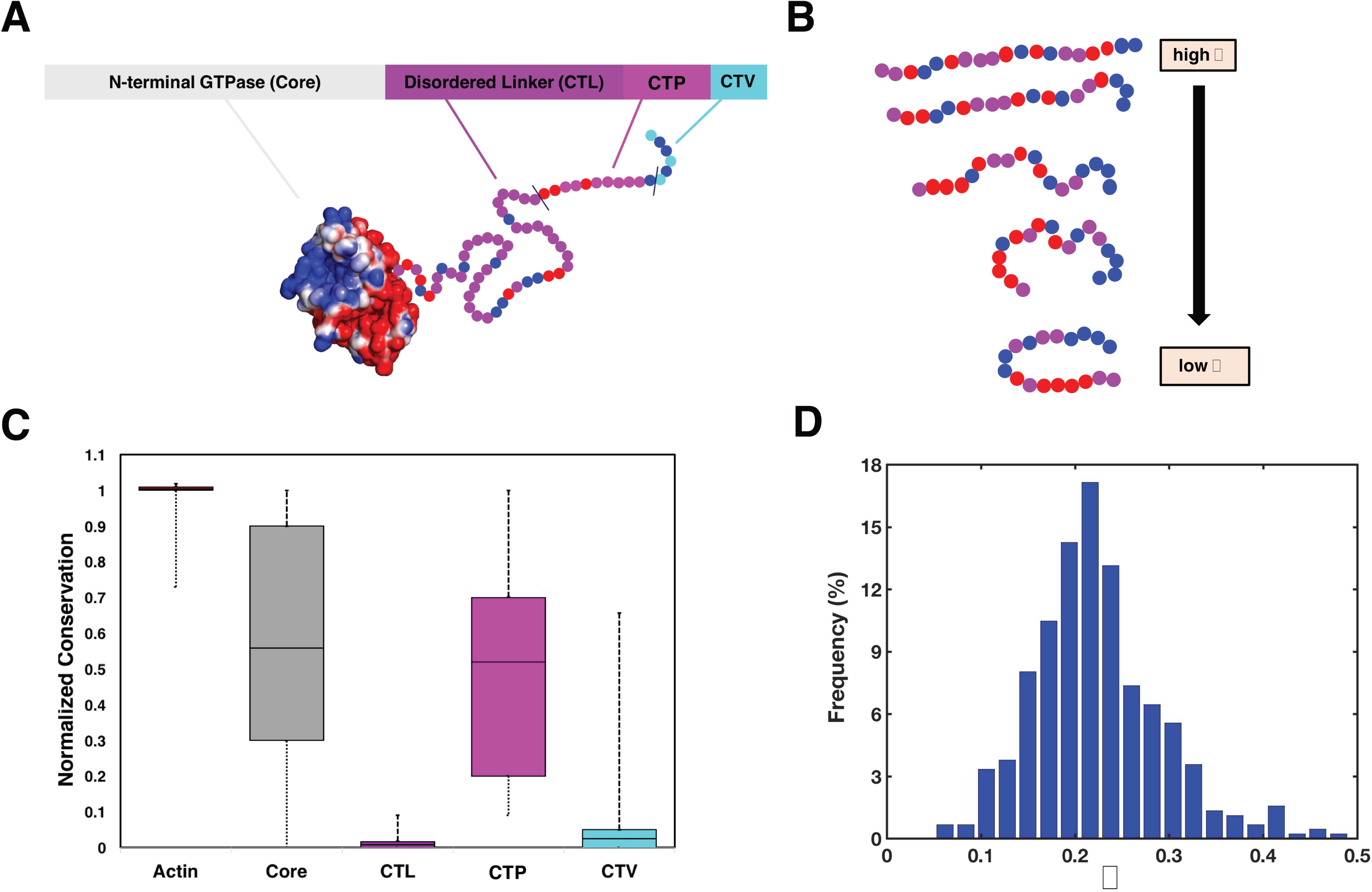
Sequence features of FtsZ from rod-shaped bacteria. (A) The domain architecture of FtsZ is characterized by an N-terminal folded core, which contributes to the GTPase activity, an intrinsically disordered C-terminal linker (CTL), a conserved 17-residue C-terminal peptide (CTP), and a C-terminal tip known as the C-terminal variable (CTV) region. The core is shown using an electrostatic surface representation that was generated using the APBS software package (Jurrus et al., 2018). Regions of high negative and positive potential are shown in red and blue, respectively; and regions that are net neutral are shown in white. The potentials are contoured in units of *k*_B_*T/e*, where *k*_B_*T* denotes thermal energy and *e* is electronic charge. Residues of the CTL, CTP, and CTV are shown as beads, with blue, red, and purple beads correspondingly denoting positively charged, negatively charged, and neutral residues. (B) For a polyampholytic IDR, the sequence-encoded κ value, which quantifies the degree of charge segregation, will fall between 0 and 1. The value of is small and approaches zero for well mixed and increases, approaching unity for sequences where the oppositely charged residues are segregated within the linear sequence. The screening of intra-chain repulsions by attractions leads to expanded conformations low κ sequences. As κ increases, attractions between oppositely charged blocks induce chain compaction. (C) Normalized conservation scores for different regions of FtsZ. The conservation scores, in terms of percent sequence identities, were computed using CLUSTAL-W. In each box and whisker plot, the solid horizontal line represents the mean conservation score. The box outlines the middle 50% of the values, and the whiskers represent the entire distribution. All conservation scores were normalized to the scores for sequences of G-actin. By definition, the mean normalized score is one for G-actin. The mean normalized conservation scores are ∼0.57 and ∼0.55 for the core domain and CTP motifs, respectively. This is to be contrasted with the low scores for the CTL and CTV sequences. (D) Although CTL sequences of FtsZ proteins are poorly conserved, >95% of the observed κ-values, which quantify the degree of segregation versus mixing of oppositely charged residues, lies between 0.1 and 0.4. These statistics were based on CTT sequences (CTL + CTP + CTV) derived from 1209 FtsZ orthologs.

Previous studies showed that deletion of the CTL leads to aberrant FtsZ assembly that compromises Z-ring formation and cell division (Buske and Levin, 2013). Replacing the wildtype (WT) CTL with a scrambled sequence variant of the linker preserves Z-ring formation and cell division. In contrast, replacing the CTL with a rigid alpha-helical domain from human beta-catenin yielded diffuse FtsZ puncta that compromise Z-ring formation and cell division (Buske and Levin, 2013). This work underscored the importance of the disordered CTL in FtsZ assembly and Z-ring formation. It also raised several questions regarding the contribution of the intrinsically disordered CTL to the assortment of functions coordinated by FtsZ. Here, we pursue answers to some of these questions by leveraging insights that have emerged through systematic studies of sequence-encoded conformational preferences of intrinsically disordered proteins / regions (Das et al., 2015).

We focus here on the CTL of *B*. *subtilis* FtsZ because its amino acid compositional bias is similar, in terms of the fraction of charged residues and net charge per residue, to that of a third of the CTL sequences in rod-shaped bacteria (Buske et al., 2015). The fraction of charged residues is 0.33 and the net charge per residue is +0.06 implying that the CTT is polyampholytic. The overall sizes, shapes, and amplitudes of conformational fluctuations of polyampholytic sequences are governed by a combination of the fraction of charged residues, the net charge per residue, the length of the IDR, and the linear patterning of oppositely charged residues (Das et al., 2015). In polyampholytic IDRs, oppositely charged residues can be separated into distinct blocks or they can be uniformly mixed along the sequence (Figure 1B). The linear mixing/ segregation of oppositely charged residues can be quantified using different parameters (Das and Pappu, 2013; Sawle and Ghosh, 2015; Sawle et al., 2017). The one we use here is designated as κ, where 0 ≤ κ ≤ 1 (Das and Pappu, 2013). If oppositely charged residues are well mixed, then the sequence will have low κ-values. In contrast, separation of oppositely charged residues into discrete blocks within the linear sequence leads to higher values of κ. In well-mixed sequences, preferential solvation of charged residues combined with a counterbalancing of intra-chain electrostatic repulsions and attractions will promote chain expansion. Increased linear segregation of oppositely charged residues causes systematic compaction of IDRs leading to smaller values for radii of gyration, more spherical shapes for individual molecules, and decreased amplitudes of global conformational fluctuations (Figure 1B).

A recent analysis of polyampholytic IDRs drawn from the human proteome showed an inverse correlation between median κ-values and median values for the fraction of charged residues (Holehouse et al., 2017). Importantly, based on random distributions of disordered sequences with an FCR ≈ 0.3 there is a clear evolutionary preference for sequences with higher than randomly expected κ-values (Figure S1). Interestingly, previous studies also showed that there is a clear selection against sequences with higher values of κ (> 0.45) (Holehouse et al., 2017). These observations also apply to CTT sequences drawn from 1209 orthologous FtsZ proteins (Buske et al., 2015). Here, we explore the functional and phenotypic impacts of charge segregation / mixing within the polyampholytic CTT of FtsZ in *B*. *subtilis*. We studied the impact of varying κ within the CTT of *B*. *subtilis* FtsZ and quantified the impact of changing κ on CTT conformations, cell division, Z-ring formation, cell growth, and the biophysical aspects of FtsZ polymerization and GTPase activity.

## Results

### CTT κ-values span a narrow range even though CTL sequences are hypervariable

We quantified the evolutionary conservation of different regions within FtsZ across 1209 orthologous proteins. We used the sequences of G-actin derived from orthologs as a reference for this analysis because these sequences are highly conserved across evolution. We used the mean conservation score for G-actin to normalize the conservation scores for different regions of FtsZ. The results are shown in Figure 1C. The core of FtsZ and the CTP motif show considerably higher degrees of conservation when compared to the CTL and CTV regions, which are poorly conserved.

We also calculated the distribution of κ-values for CTT sequences derived from 1209 distinct orthologs (Figure 1D). Given the conservation of the CTP, it follows that the variation in κ is mainly due to sequence variations within the CTL. Roughly 95% of the CTT κ values are within the interval 0.13 < κ < 0.36. The CTT from *B*. *subtilis* FtsZ has a κ value of 0.19. The expected distribution of κ values for randomly shuffled sequences that are constrained to have the CTT composition of *B*. *subtilis* FtsZ is shifted to lower values when compared to the actual κ values observed for the 1209 orthologous sequences (Figure S2). However, when compared to the shuffled sequences generated in the absence of any selective pressure, many sequences with higher κ values are found, although very few naturally occurring sequences have a κ value above 0.4. This suggests that there is an evolutionary pressure against CTT sequences with κ values that go above 0.4. Similarly, there appears to be a selection pressure against sequences with κ values lower than 0.15.

### Design of sequence variants of *B*. *subtilis* FtsZ

To test the impact of changing CTT κ values on FtsZ function, we designed a set of sequence variants of *B*. *subtilis* FtsZ. In these variants, we fixed the GTPase core domain, the amino acid composition of the CTL, and the sequence of the CTP. We shuffled the positions of charged and neutral residues within the *B*. *subtilis* CTL to generate sequence variants of the CTT that span the spectrum of κ values between 0.1 and 0.8. There are roughly 10^40^ sequences that satisfy the design criteria (Figure 2A). Of these, we selected six variants of *B*. *subtilis* FtsZ and each of these variants is distinguished by the κ value for the CTT, which spans the range from 0.15 to 0.72 (Figure 2A and Figure 2B). Each of the designed *B*. *subtilis* FtsZ variants is designated as CTLV*x*, where *x* is an integer value that increases as the CTT κ value increases.

**Figure 2:**
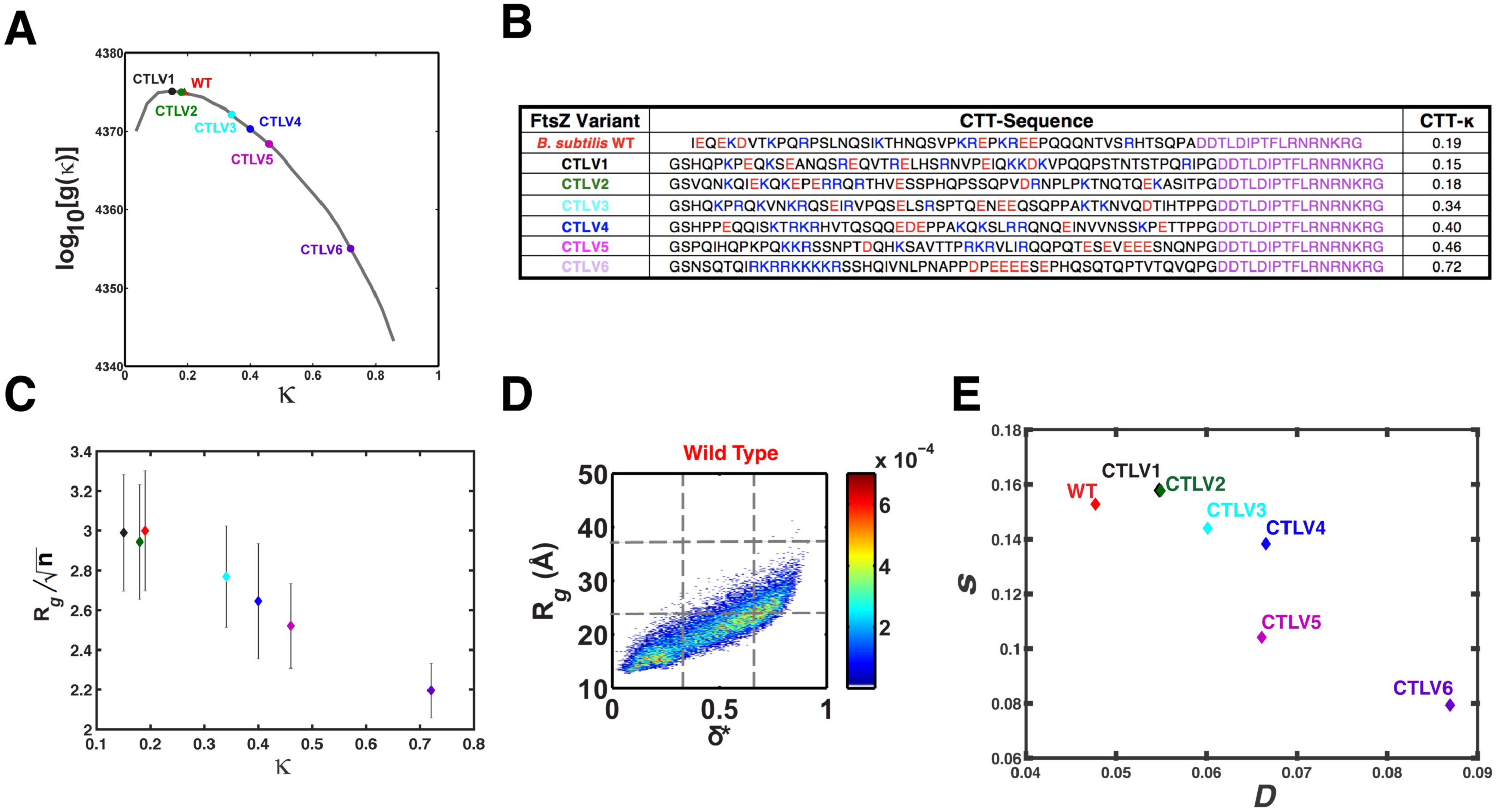
Designed CTT sequences and their global conformational properties. (A) Keeping the CTP sequence fixed, we designed sequence variants of the *B. subtilis* FtsZ by fixing the amino acid compositions and shuffling residues within the CTL to generate proteins with CTT sequences that have altered linear patterning of oppositely charged residues. There are roughly 10^40^ sequences that meet the design criteria. (B) We chose six CTT sequences with κ values spanning from 0.15 to 0.72. The CTT sequences for each FtsZ variant are shown in the table in panel (B). Column 1 shows the name that was assigned to each *B. subtilis* FtsZ variant; columns 2 and 3 show the redesigned sequence of the CTL and the κ-value for the variant in question, respectively. Values of κ were calculated over the entire CTT, and this includes the CTL and CTP sequences. (C) Plots of length normalized values of the mean radii of gyration for each of the designed CTT variants. These values were obtained by dividing the mean *R_g_* values for each variant by the square root of the length *n* of the corresponding sequence. The extra residues present in each CTL variant were a product of cloning. These data show that the length normalized *R_g_* is high for CTT sequences with κ values that are low. In contrast, we observe a monotonic decrease as κ values increase beyond that of the WT. (D) Two-dimensional conformational distributions from atomistic simulations of CTT sequences for the WT. These distributions quantify the joint probability density *p*(*R_g_*,δ^*^) of sizes (R_g_) and shapes (δ^*^). Results for the six designed CTT variants are shown in Figure S2A. In the simulations, each CTT variant was modeled as an autonomous unit. Each sub-panel in Figure S2A shows the sequence-specific two-dimensional histogram of shapes, quantified in terms of asphericity values (δ^*^) and sizes calculated in terms of radii of gyration (*R_g_*). In each sub-panel, the abscissa is divided into nine equal intervals with the boundaries δ^*^ < 0.33, 0.33 < δ^*^ ≤ 0.66, δ^*^ > 0.66 and 10 Å ≤ *R_g_* ≤ 23 Å, 23 Å ≤ *R_g_* < 36 Å, and 36 ≤ ≤ *R_g_* ≤ 50 Å. (E) Plot of the Shann entropy (*s*) versus *D* for the WT CTT and each of the designed CTT variants. The Shannon entropy *s* for each sequence variant is calculated as: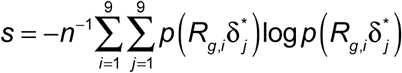where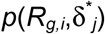is the probability density associated with bin (*i*,*j*) on the sequence-specific twodimensional histogram (panel **D** for the WT CTT and Figure S2A for each of the six variants).

### Sequence-intrinsic conformational preferences of CTT sequences are influenced by the extent of charge segregation

We performed atomistic simulations of the WT CTT from *B*. *subtilis* FtsZ and the six designed variants. We used the ABSINTH implicit solvation model and forcefield paradigm (Vitalis and Pappu, 2009a; Vitalis and Pappu, 2009b) for these simulations. The results were used to quantify the impact of charge patterning on the overall sizes, shapes, and amplitudes of conformational fluctuations of CTT sequences modeled as autonomous units. As κ increases, there is a clear trend toward compaction. We quantified the preference for compact structures in terms of the length normalized mean radii of gyration 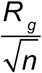
that are plotted for the WT CTT and designed variants (Figure 2C). For each variant, 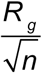 is calculated as the mean *R_g_* value of the variant divided by the square root of the number of residues *n* in the CTT. Variants of the *B*. *subtilis* CTT sequence with κ values that are lower than the WT are slightly more expanded than the WT CTT, although the degree of expansion is not statistically significant. However, sequences with κ values that are higher than the WT become increasingly more compact with increasing κ. It follows, in accord with previous observations for other systems (Das et al., 2016; Das and Pappu, 2013; Sherry et al., 2017),that increasing the linear segregation of oppositely charged residues strengthens intra-CTT electrostatic attractions.

The sequence-intrinsic conformational heterogeneity of the WT *B*. *subtilis* FtsZ CTT sequence is borne out in terms of the broad spectrum of sizes and shapes that this sequence adopts. These parameters are quantified in terms of the radii of gyration (*R_g_*) and asphericities (δ^*^) that are achievable via spontaneous conformational fluctuations (Figure 2D). Distributions of *R_g_* and δ^*^ for each of the designed CTT sequences are shown in Figure S3A. The diversity of sizes and shapes can be quantified using the (*R_g_*, δ^*^) distributions and summarized in terms of a normalized Shannon entropy (*s*). Larger values of *s* imply a greater diversity of and sizes and shapes.

A large number of distinct conformations might be compatible with similar values of *R_g_* and δ^*^. Here, we refer to this feature as the ruggedness of the energy landscape. Following previous approaches (Lyle et al., 2013), we quantify ruggedness in terms of the degree of similarity / dissimilarity of backbone dihedral angles among every pair of conformations sampled by each variant. This yields a distribution of similarity / dissimilarity values denoted as *p*(*D*) and we take the mean value, denoted as *D* to be an average measure of backbone dihedral angle similarity / dissimilarity. For a well-folded protein, such as the FtsZ core, the value of *D* will be small because the backbone dihedral angles of distinct conformations that are accessible as a result of spontaneous fluctuations will be similar to one another. In contrast, the value of *D* will be higher for systems that sample conformations with distinctly different patterns of backbone dihedral angles.

The combination of values for *s* and *D* will define the sequence-specific topography of the energy landscape. This aspect is schematized in Figure S3B in terms of a one-dimensional projection of the energy landscape along a single conformational coordinate that combines the contributions of sizes, shapes, and backbone dihedral angles. Landscapes can be narrow and smooth (low *s* and *D*), narrow and rugged (low *s* and large *D*), wide and smooth (high *s* and low *D*), or wide and rugged (high *s* and *D*). Figure 2E shows the values of *D* and *s* for the WT CTT and each of the designed CTT variants. These results show that of the seven CTT sequences, the WT sequence best conforms to the designation of encoding a wide and smooth landscape. The energy landscapes of CTT sequences become progressively more rugged (increasing *D*) with increasing κ, although the landscapes for variants CTLV1, CTLV2, CTLV3, and CTLV4 retain the width (similar *s* values) of the WT CTT. The energy landscapes of CTT sequences from CTLV5 and CLTV6 are fundamentally different in that they become narrower (smaller *s* values) and more rugged (larger *D* values). Overall, our analysis shows that the WT CTT stands out as the sequence with nearly the highest conformational diversity in terms of global sizes and shapes (high *s* value) and the lowest ruggedness, given that it has the lowest *D* value.

### Cellular phenotypes are robust in *B*. *subtilis* that express variants CTLV2, CTLV3, and CTLV4

We used in cell investigations to probe the impact of designed changes to CTT sequences on cell division and cell growth in *B*. *subtilis*. Western blots in Figure 3A show that the cellular levels of FtsZ variants are sensitive to sequence-encoded features within the CTT. FtsZ variants with extreme κ values for their CTT sequences such as CTLV1 and CTLV6 are not well tolerated by the cells (Figure 3A). This in turn influences Z-ring formation because it diminishes the cellular levels of the FtsZ variants. Clone verification using restriction digests and sequencing at the vector-insert junctions showed that the constructs were correctly cloned. Instead, it appears that variants that have low cellular levels are actively degraded as opposed to being under expressed.

**Figure 3:**
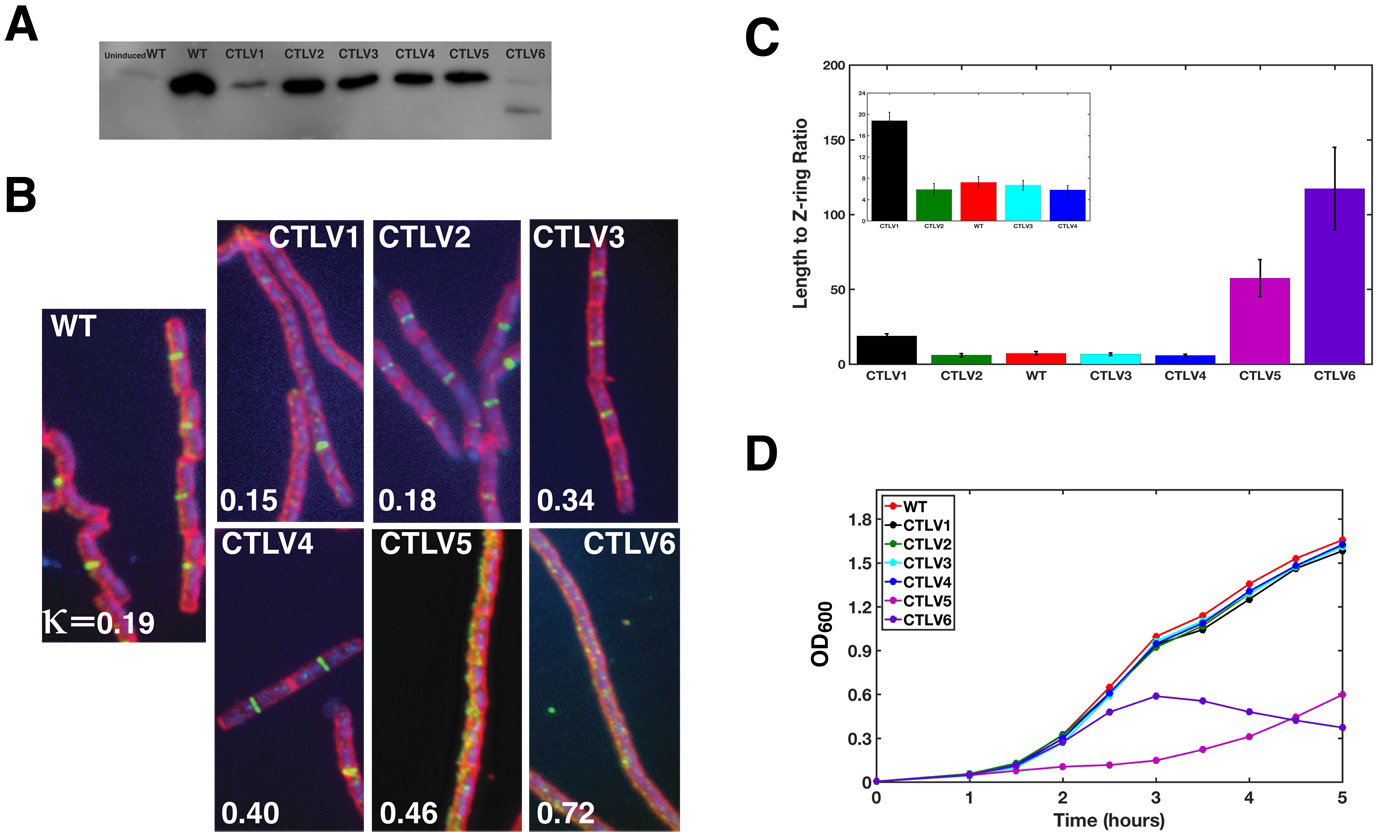
Phenotypic impact of changing the CTT κ-values with fixed amino acid composition and an invariant CTP. (A) Western blot quantifying the protein levels of each variant, normalized to the WT FtsZ, in *B. subtilis.* The concentration of CTLV1 is much lower than that of WT FtsZ, suggesting that sequences with lower than WT κ-values are prone to degradation. CTLV6 shows two distinct bands. All other variants are expressed and retained to levels that are comparable to that of WT FtsZ. (B) Immunofluorescence micrographs of *B. subtilis* expressing the WT FtsZ (left) versus six different variants of FtsZ with redesigned CTL sequences (shown on the right as a 2×3 grid of micrographs). Each micrograph includes the label of the variant (top) and the κ-value of the CTT sequence (bottom). Z-ring formation is robust for variants CTLV2-CTLV4. In contrast, for variants with CTT κ-values that are larger than 0.40 (see CTLV5 and CTLV6), we observe a haphazard distribution of puncta as opposed to robust Z-ring formation. For CTLV1, we observed both properly formed Z-rings and a diffuse distribution of FtsZ. (C) Length-to-Z-ring (L/R) ratio for *B. subtilis* expressing different FtsZ variants. For the WT FtsZ, this ratio is ∼5. The value of L/R is similar to that of WT for cells expressing variants CTLV2, CTLV3, and CTLV4. The error bars quantify the standard error in our estimate of the mean L/R values. (D) *B. subtilis* expressing FtsZ variants CTLV5 and CTLV6 (κ > 0.4) show a growth phenotype in contrast to all other variants that have WT profiles for growth. Data for growth phenotypes are shown in terms of the normalized optical density at 600 nm (OD_600_). This quantifies the normalized concentration of viable bacteria in a suspension.

In cells where CTLV1 reaches a sufficient concentration, well-defined Z-rings are formed (Figure 3B). The cellular levels of CTLV2, CTLV3, and CTLV4 are not compromised by the designed changes to the CTT sequences (Figure 3A). Cells expressing these three variants show robust Z-ring formation. This is true even though the κ values for their CTT sequences span a range from 0.18 – 0.4. However, Z-ring formation is compromised for cells expressing variants CTLV5 (κ=0.46) and CTLV6 (κ=0.72) (Figure 3B). These results suggest that increased segregation of oppositely charged residues within the CTT sequence leads to a disruption of Z-ring formation and haphazard localization of FtsZ puncta that in turn inhibit cell division.

To quantify the robustness / disruption of Z-ring formation, we measured the L/R ratio, which describes the ratio of the average length of the cell to the number of Z-rings observed (Buske and Levin, 2013). This is a measure of the fitness and robustness of cell division in rodshaped bacteria. The L/R ratio is 7.2 ± 1.2 for cells expressing WT FtsZ. Compromised fitness is indicated by L/R ratios that are larger than the WT. The L/R ratio is 5.9 ± 1.1, 6.6 ± 1.0, and 5.8 ± 0.9 for cells expressing variants CTLV2, CTLV3, and CTLV4, respectively (Figure 3C). These measurements suggest robust Z-ring formation and division for cells expressing FtsZ variants with CTT κ values between 0.18 and 0.40. In contrast, the L/R ratio is 18.8 ± 1.6, 57.5 ± 12.5, and 117.3 ± 27.6 for cells expressing variants CTLV1 (CTT κ=0.15), CTLV5 (CTT κ=0.46), and CTLV6 (CTT κ=0.72), respectively.

The higher L/R ratio for CTLV1 is likely due to lowered cellular levels (Figure 3A). The *in vivo* behavior we observe for cells expressing CTLV5 and CTLV6 cannot be explained in terms of reduced protein levels alone (Figure 3B). Therefore, as an additional measure of cell division efficiency we monitored the rate of change of optical density at 600 nm (OD_600_), which is proportional to the concentration of cells in mid-log phase of growth. A reduction in OD_600_ would arise from the combination of a decrease in the efficiency of cell division, compromised metabolism, and increased cell death. As shown in Figure 3E, cells expressing CTLV5 and CTLV6 show a reduction in the rate of increase of OD_600_, indicating that CTLV5 and CTLV6 cause a significant reduction of the cellular growth rate. In contrast, cells expressing all other variants, including CTLV1, showed growth rates that are similar to those of cells expressing WT FtsZ.

### *In vitro* polymerization, bundling, and GTPase activity of CTLV4 is comparable to that of WT

The GTP binding site and the T7 activation loop that is required for the catalytic activity are on opposite sides of the core domain of FtsZ. Accordingly, the active form of the enzyme is a dimer and GTP is hydrolyzed at the interface of the GTP binding site and the T7 loop (Figure S4A). GTP binding promotes dimerization and the formation of higher-order head-to-tail polymers of FtsZ. In the dimeric state, GTP hydrolysis at the subunit interface leads to dissociation into two monomers (Scheffers et al., 2002). In the polymeric form, GTP hydrolysis promotes bending and ring formation (Erickson et al., 2010).

Recent studies have shown that the CTT also has a role to play in polymerization and Z-ring formation, *in vitro* and *in vivo* (Buske and Levin, 2013). Right-angle light scattering is a sensitive method for studying FtsZ polymerization and bundling of FtsZ polymers in the presence of GTP. Outputs from a typical light scattering assay are shown in Figure S4B. In these experiments, the concentration of FtsZ is above the apparent critical threshold for polymerization, which was found to be 2.5 μM in the presence of 50 mM KCl, in a pH 6.5, 50 mM MES buffer (Caplan and Erickson, 2003). Upon the introduction of 1 mM GTP into the solution mixture, the scattering intensity increases within a narrow time interval and rapidly reaches a steady state value. Light scattering is sensitive to the presence of large aggregates and these might include linear polymers, bundled polymers, or even large spherical assemblies. We collected samples at the mid-point of the scattering assay and performed deep etch electron microscopy imaging. A representative image for WT FtsZ is shown in Figure 4A. These images reveal the formation of linear and curved protofilaments that are roughly 100 nm long.

**Figure 4:**
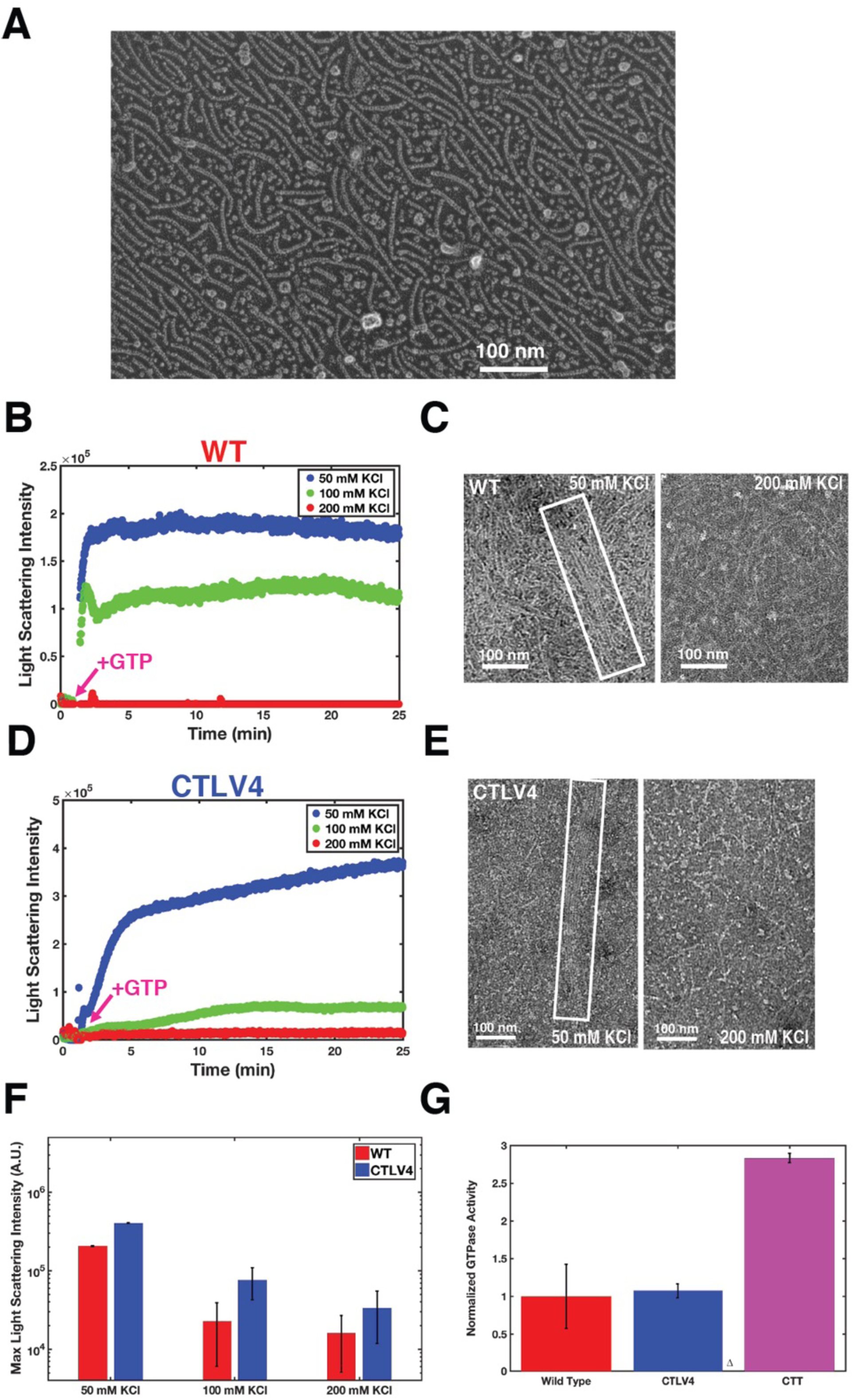
FtsZ forms bundled filaments. (A) Representative deep etch electron microscopy image of protofilaments formed by the WT FtsZ in the presence of GTP. (B) Representative data traces from light scattering for WT FtsZ in the presence of differing amounts of KCl. The arrow marks the time point for introduction of GTP into the solution mixture. At low salt, linear FtsZ filaments form bundled assemblies. The bundling is weakened at higher salt concentrations. (C) The differences in morphologies, specifically with regard to the extent of bundling of filaments, are shown in electron micrographs obtained in 50 mM KCl versus 200 mM KCl. Based on these morphological data, we propose that the light scattering data are mainly sensitive to the extent of bundling of filaments. (D) Analogous light scattering data for CTLV4. (E) Negative stain electron microscopy images showing the morphologies formed by assemblies of CTLV4 molecules in 50 mM KCl versus 200 mM KCl. In low salt, the CTLV4 molecules make long, bundled polymers. These polymers shrink in length and become unbundled at high salt. (F) Comparison of the extent of bundling as a function of salt. This is quantified in terms of the normalized maxima in the scattering profiles for the WT and CTLV4 variant in the presence of different amounts of KCl. For WT and CTLV4, bundling decreases as salt concentration increases. (G) Comparative assessments of GTPase activity – normalized to the WT FtsZ – for CTLV4 and ΔCTT. Polymerization weakens the GTPase activity and this auto-inhibition derives from the presence of the tail. This point is made by comparing the efficiencies of GTP hydrolysis for CTLV4 and ΔCTT to that of the WT FtsZ. The efficiencies are shown normalized to that of WT. Absence of the CTT leads to an enzyme that almost three times more efficient as a GTPase when compared to the WT or CTLV4.

The light scattering data in Figure S4B were obtained in the presence of 50 mM KCl. Previous studies showed that FtsZ can form bundled polymers and the extent of bundling is dependent on salt concentration (Buske and Levin, 2012; Erickson et al., 2010). Figure 4B shows comparative light scattering traces obtained in the presence of different concentrations of KCl. The steady-state scattering intensity decreases sharply as salt concentration is increased. Negative stain EM imaging shows that the FtsZ polymers form large bundles at low salt and this bundling is disrupted at high salt (Figure 4C). The EM data together with the light scattering data suggest that the scattering intensity derives mainly from the bundling of FtsZ polymers, as opposed to the formation of linear polymers.

Next, we studied the salt dependence of the light scattering traces obtained for CTLV4. This variant is of interest because it maintains cellular phenotypes despite an increase of the CTT κ by a factor of two when compared to the WT CTT. Like WT, we observed a canonical dependence of light scattering intensity on monovalent salt (Figure 4D). At high salt, the interactions that hold the bundles together are screened and CTLV4 forms unbundled polymers that are similar to those formed by WT FtsZ (Figure 4E). Simulations predict the CTT of CTLV4 is 10% smaller than the WT CTT (Figure 2C). Compaction is mediated by increased electrostatic interactions between charged blocks on the CTT sequence. While the simulations illustrate charge-dependent intramolecular interactions, these interactions could also occur in an intermolecular fashion. Concordant with this expectation, we observed a two-fold increase in the peak scattering intensity in the presence of 50 mM KCl for the CTLV4 variant when compared to the WT FtsZ (Figure 4F). This increase in scattering intensity derives from two distinct features as revealed by negative stain EM images (Figure 4E). When compared to the WT FtsZ, CTLV4 forms longer polymers that become part of thicker bundles in the presence of low salt. Figure 4F shows a comparison of the relative strengths of filament / polymer bundling by comparing the maximum scattering intensities measured for the WT versus CTLV4 in the presence of three different concentrations of KCl. The major differences between the variants are manifest at low salt. However, under physiologically relevant salt concentrations, the morphological differences between CTLV4 and WT FtsZ are minimal. Therefore, while the CTT of CTLV4 can be involved in enhanced tail-mediated interactions, especially at low salt, this does not deleteriously affect polymerization or bundling of CTLV4. These data provide an explanation for the observed robustness of Z-ring formation and cell division for cells expressing CTLV4 as well as cells expressing variants CTLV2 and CTLV3.

To further assess the robustness of altering charge patterning within evolutionarily observed bounds (Figure 1D), we quantified the comparative efficiencies of GTP hydrolysis of FtsZ variants. Figure 4G shows results from measurements that compare the efficiencies of WT FtsZ to those for CTLV4 and ΔCTT. The latter is a variant of FtsZ that lacks the CTT. The efficiency of GTP hydrolysis remains unchanged for CTLV4 when compared to the WT FtsZ (Figure 4G). Higher-order assembly, such as protofilament bundling and Z-ring formation, are compromised for ΔCTT (Buske and Levin, 2013; Sundararajan and Goley, 2017; Sundararajan et al., 2015) and we do not observe a measurable increase in light scattering that is distinguishable from the background. Despite this, we find that ΔCTT is much more efficient as a GTPase with an enzymatic activity that is ∼3 times higher than the WT or CTLV4. Taken together with the observation that GTP hydrolysis promotes the severing of protofilaments (Mukherjee and Lutkenhaus, 1998) our data suggest that ΔCTT, as a result of being unencumbered by the absence of the CTT, is able to bind GTP, polymerize, hydrolyze GTP, disassemble and repeat the cycle in an extremely efficient manner. Accordingly, we propose that the CTT, which is essential for Z-ring formation, promotes FtsZ assembly and this suppresses GTPase activity. This suppression could arise from steric effects of the CTT, a reduction in subunit turnover, and possible protofilament stabilization through interactions with neighboring filaments mediated by the CTT.

### CTLV5 and CTLV6 drive alternative modes of polymerization even in the absence of GTP

Given the deleterious phenotypic impacts associated with CTLV5 and CTLV6, we asked if these variants promote or hinder FtsZ polymerization *in vitro*. We performed light scattering assays for CTLV5 and CTLV6 in the presence and absence of GTP. The light scattering data obtained in the presence of GTP show a WT-like decrease in light scattering intensity at physiological concentrations of monovalent salt (Figure 5A). In 50 mM KCl, CTLV5 forms large linear filaments that are not bundled (Figure 5B). Surprisingly, even in the absence of GTP, CTLV5 forms discernible assemblies that persist at high salt concentrations (Figure 5C). This behavior is distinct from what we observe for WT FtsZ and CTLV4. The assemblies formed by CTLV5 in the absence of GTP are spherical aggregates that are roughly 20 nm in diameter (Figure 5D). Interestingly, light scattering from these assemblies that form in the absence of GTP, increases with increased salt concentration. Relative CTLV5 light scattering magnitudes for different buffer conditions are shown in Figure 5E. The increased assembly size in high salt might derive from a weakening of electrostatic repulsions amongst the surfaces of assemblies. This is clearly a CTT-mediated phenomenon given that we do not observe these assemblies for the wild-type FtsZ or for variants with lower degrees of charge segregation within their CTTs.

**Figure 5:**
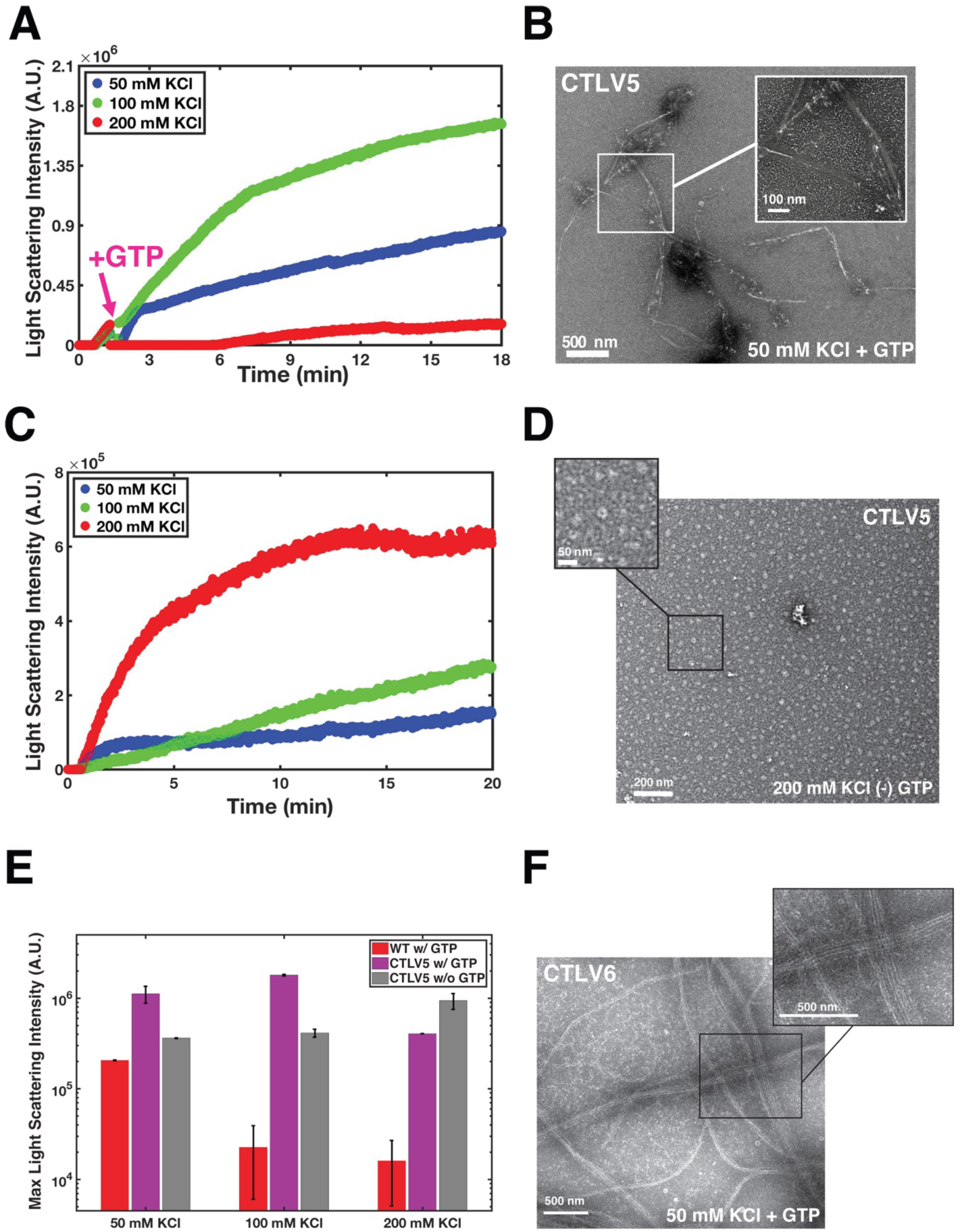
Increased charge segregation within CTT sequences leads to tail-mediated assembly and diminution of GTPase activity. (A) Salt dependence of light scattering profiles for CTLV5 in the presence of GTP. (B) Negative stain EM images show that in the presence of GTP, the CTLV5 variant of FtsZ forms canonical polymers that are also highly bundled. The bundling is weakened in high salt (200 mM KCl). (C) In the absence of GTP, we observe significant light scattering for CTLV5 variants. The scattering intensity increases in the presence of higher concentrations of KCl. (D) Negative stain EM images show that CTLV5 molecules form large, spherical aggregates in the absence of GTP. (E) Comparative assessments of the maxima in light scattering for CTLV5 with and without GTP referenced to that of WT FtsZ in the presence of GTP. As salt concentration increases, GTP-dependent assembly is weakened. However, an increase in salt concentration facilitates the formation of the larger spherical aggregates for CTLV5. (F) Negative stain EM images of morphologies obtained for CTLV6. Increased segregation of oppositely charged residues leads to the formation of linear filamentous “tracks” characterized by inter-filament interactions involving the CTTs as shown in the EM image.

Of the designed variants, CTLV6 has the most extreme degree of segregation of oppositely charged residues in its CTT. Increased segregation of oppositely charged residues within the CTT drives increased self-association of FtsZ via inter-tail interactions between filaments. These interactions generate profoundly different assemblies when compared to the morphologies that we observe for wild-type FtsZ or other designed variants (Figure 5F). The spacing between filaments corresponds to the length of a fully extended CTT. This is suggestive of zippering interactions between tails with “blocky” (charge-segregated) sequences. These assemblies persist at physiological salt concentrations in the presence of GTP (Figure S5A & B), indicating that they likely form in the cellular setting as well. At 200 mM KCl, we observe morphologies that are similar to those seen in 50 mM KCl (Figure S5C). We further observed a markedly lower rate of GTP hydrolysis for CTLV6 when compared to that of WT. This is indicative of reduced rates of subunit turnover for assemblies formed by FtsZ variants with high κ CTT sequences (Figure S5D).

### Delineating the contributions of charge patterning and amino acid composition within CTT sequences

The CTT sequences of all of the designed variants studied to this point have identical amino acid compositions that were derived from the CTT of *B*. *subtilis* FtsZ. The parameter κ quantifies the extent of segregation or mixing of oppositely charged residues within the CTT and is agnostic to the specific amino acid composition. In order to separate the contributions from amino acid composition and charge patterning, we designed an FtsZ variant with a WT-like κ for its CTT. However, the amino acid composition was based on a reduced amino acid alphabet. The composition of the CTT was such that all positive charges from the CTL of *B*. *subtilis* FtsZ were replaced with Arg, all negatively charged residues were replaced with Glu, Ser / Thr / Cys residues were replaced with Ser, all Asn / Gln residues were replaced with Gln, all hydrophobic groups were replaced with Ala / Val, and the numbers of Gly / Pro residues were kept intact (Figure 6A). The FtsZ variant with a reduced alphabet and κ=0.21 CTT is tolerated in *B*. *subtilis*. This variant, designated as RLWT forms robust Z-rings and grows in a wild-type manner (Figure 6B, 6C, & 6D). However, the cellular levels were significantly lowered for FtsZ variants with reduced alphabet for their CTT sequences and κ values of 0.17 (RLV1) and 0.47 (RLV5), so designated because they were intended to be mimics of CTLV1 and CTLV5 (Figure 6B). In these cells, Z-ring formation and cellular growth are also impaired (Figure 6C & 6D).

**Figure 6:**
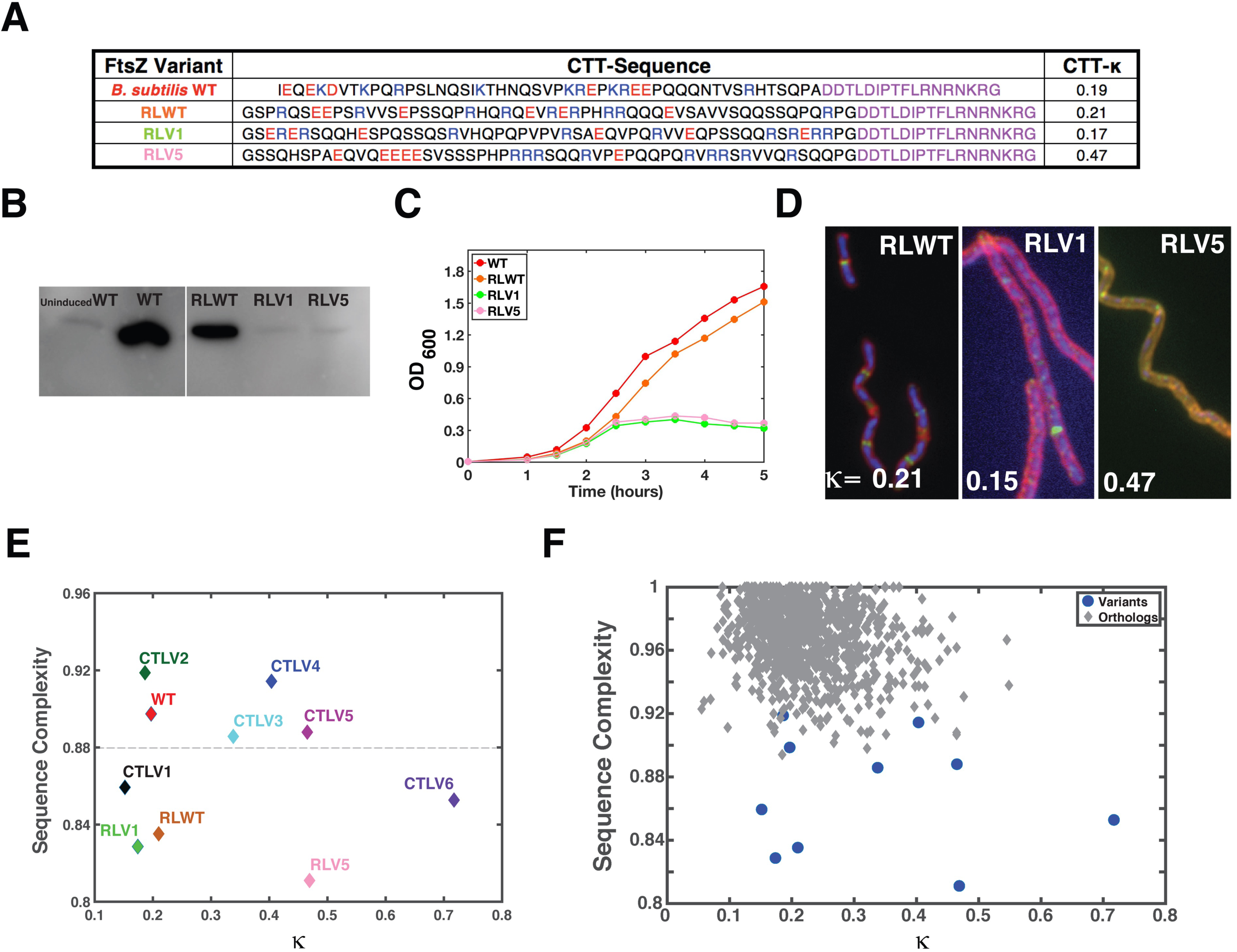
Dissecting the contributions of κ, amino acid composition, and sequence complexity on FtsZ assembly and cell division. (A) Designed CTT sequences based on a reduced amino acid alphabet. The corresponding FtsZ variants are referred to as RLWT, RLV1, and RLV5 because the CTT κ values of these variants are similar to those of WT, CTLV1, and CTLV6. Here, RL implies reduced library. Column 3 shows the κ values for each of the CTT sequences. (B) Immunoblot analysis of the cellular levels of RLWT, RLV1, and RLV5 compared to the WT FtsZ. The lower levels of RLV1 and RLV5 suggest that, in *B. subtilis*, these variants are degraded more readily than WT FtsZ or even RLWT. (C) Comparative growth curves for B. subtilis expressing WT FtsZ, RLWT, RLV1, and RLV6, respectively. Cells expressing RLV1 and RLV6 show a dominant negative growth phenotype, mirroring the behavior of cells expressing CTLV5 and CTLV6. (D) Immunofluorescence micrographs showing the robust Z-ring formation for RLWT and poor localization and haphazard distributions of RLV1 and RLV5, respectively. (E) Plot of the LZ sequence complexity (see main text) (Lempel and Ziv, 1976) of each of the nine CTT sequences including those from WT FtsZ and the eight designed variants. Calculations of the LZ sequence complexities were performed using localCider (Holehouse et al., 2017) and the results were plotted against the variant-specific κ values. The plot shows that the LZ values are greater than 0.88 for the CTT of WT FtsZ and the CTT sequences of CTLV2, CTLV3, CTLV4, and CTLV5. The LZ values fall below a threshold of 0.88 for the CTT sequences of variants CTLV1, CTLV6, RLWT, RLV1, and RLV5. These calculations show that variants with low protein stability have CTT sequences with LZ values that are below a threshold value of 0.88. (F) Plot of LZ complexity values for CTT sequences from 1209 FtsZ orthologs (gray diamonds). All of the CTT sequences have LZ complexity values that lie above the threshold value of 0.88 with an overwhelming number of sequences having values greater than 0.92. This suggests an evolutionary selection for CTT sequences that encode a combination of higher LZ values and optimal κ values.

### Cellular levels of FtsZ variants are lowered for those with reduced complexity in their CTT sequences

Measures of sequence complexity quantify the diversity of the amino acid alphabet and the complexity of the sequence syntax. For sequences of similar amino acid compositions, the sequences that cluster similar amino acids into contiguous blocks will have lowered sequence complexity. Sequence complexity is also lowered by using a simplified amino acid composition as we have done with the CTT sequences that are based on a reduced amino acid library. In eukaryotic systems, ubiquitinated substrates engage productively with the proteasome if and only if they have low sequence complexity tags at their termini (Kraut et al., 2012; Schrader et al., 2011). Accordingly, we quantified the sequence complexities of each of the designed CTTs based on the WT and reduced amino acid alphabets. Here, we use the Lempel-Ziv (LZ) measure of complexity (Lempel and Ziv, 1976) that has been used in previous analysis of IDPs / IDRs (Holehouse et al., 2017; Romero et al., 2000). Interestingly, we find that the variants that have low cellular levels in *B*. *subtilis* (Figure 3B & 6B) also have the lowest LZ sequence complexity in their CTT sequences (Figure 6E). The sequence complexity of CTT sequences is lowered either by increasing or decreasing κ within the CTT, for the WT amino acid composition or by using a reduced alphabet for the amino acid composition.

Our analysis suggests that there is probably an evolutionary selection against FtsZ molecules with low complexity CTT sequences. This selection pressure would appear to protect against lowered cellular stability of FtsZ and tail-mediated interactions that compromise FtsZ assembly. In accord with this proposal, the LZ sequence complexity values of CTT sequences from all 1209 orthologs are higher than those of CTT sequences that engender compromised cellular stability or deleterious tail-mediated interactions (Figure 6F). These data lead to the hypothesis that a viable CTT sequence of FtsZ must have a CTL of sufficient amino acid complexity in order to avoid degradation, and an overall κ value to ensure that the tail engages in an optimal balance of intra- and inter-molecular interactions that stabilize the polymer without compromising subunit turnover kinetics or engaging in tail-to-tail associations.

## Discussion

Intrinsically disordered regions come in different sequence flavors (Das et al., 2015). However, uncovering the connections between sequence-to-conformation relationships and the molecular functions and / or cellular phenotypes that are regulated by IDRs is non-trivial. Structure-based mutagenesis experiments are not helpful because there is no singular structure that one can perturb. Instead, mutations have to be designed to shift conformational populations (Hilser and Thompson, 2007; Li et al., 2017) and fundamentally alter global and / or local conformational preferences (Das et al., 2015). One can presumably build evolutionary profiles of sequences to identify regions that appear to be highly conserved and hence immutable or covarying (Rivoire et al., 2016). Whilst such an approach would readily identify the structured regions of FtsZ, specifically the core and the CTP, as evolutionarily relevant modules, this approach is not readily transferrable to IDPs / IDRs. The CTL shows minimal sequence conservation – an attribute that is shared by many other IDPs / IDRs (Chen et al., 2006; Holehouse et al., 2017). Thus, using a sequence alignment based approach to identify relevant modules would erroneously suggest that the CTL might either be non-essential or serve as a passive tether that is interoperable with any other randomly chosen flexible sequence tether.

In order to understand the functional roles played by the CTL, we resorted to an approach driven by sequence design that is guided by physical principles underlying the sequence-to-conformation relationships of IDRs (Das and Pappu, 2013). A similar approach was pursued to uncover the roles of functional polyampholytic IDRs in p27, an inhibitor of cyclin-dependent kinases in yeast (Das et al., 2016), and the IDR within the RAM region of the Notch intracellular domain that regulates transcription of Notch genes in mammalian cells (Sherry et al., 2017).

Although the CTL sequence of FtsZ is poorly conserved across orthologs, the patterning of oppositely charged residues, as quantified by κ is bounded within a well-defined interval Figure 1D). These bounds are such that there is a pronounced rightward shift vis-à-vis a random distribution of κ values that one would calculate for a random CTL of identical amino acid composition (Figure S2). Our experiments show that the observed bounds on CTL / CTT κ values have biological and biophysical underpinnings. Additionally, variants of FtsZ with low complexity CTT sequences have low stability in the cellular milieu and are likely degraded by proteases. This presents at least a two-variable driving force for evolutionary selection whereby bacteria are likely to select against CTT sequences with low complexities and κ values that enable aberrant tail-driven assemblies, whilst selecting for sequences that enable FtsZ polymerization and controlled GTP hydrolysis / subunit turnover.

For CTT sequences with κ values that lie within the evolutionarily observed bounds, the CTTs promote lateral bundling of FtsZ filaments. These interactions are weak enough to be screened by monovalent salts such as 200 mM KCl, which is less than the basal KCl salt concentration of 350 mM in *B*. *subtilis* (Whatmore et al., 1990). For FtsZ variants with higher κ CTTs, the lateral interactions appear to have at least two distinct modes. In the presence of GTP, the tail-to-tail interactions can be weakened with monovalent salt if FtsZ molecules bind GTP and make canonical polymers. However, in the second mode of interaction, unmasked here in the absence of GTP, the salt dependence of assembly formation is inverted vis-à-vis the wild-type FtsZ, and tail-mediated interactions promote aberrant assemblies (Figure 5D & 5E). It would appear that assemblies stabilized by tail-to-tail associations can grow larger as salt concentration increases – a feature that resembles salting-out transitions whereby proteins become insoluble at higher salt concentrations (Ruckenstein and Shulgin, 2006).

The pathology of tail-mediated assemblies is made particularly vivid for CTLV6, which has a highly κ CTT (κ=0.72). The intermolecular electrostatic interactions are so strong that one observes large linear polymers with inter-polymeric “tracks” that are consistent with stabilization via tail-to-tail associations (Figure 5F). These types of strong, complimentary electrostatic interactions have also been recently reported by Borgia et al. for polyelectrolytic IDPs where oppositely charged IDPs form binary, albeit disordered high affinity complexes (Borgia et al., 2018). Electrostatic attractions can also drive phase transitions via complex coacervation as was recently demonstrated for the intrinsically disordered Nephrin intracellular domain (Pak et al., 2016). The driving forces for phase separation increase multi-fold for variants of NICD with long blocks of charged residues. Similar results were observed by altering the patterning of oppositely charged residues within the IDR of DDX4, a protein that forms nuage bodies (Nott et al., 2015). Theories show that the driving forces for phase separation become stronger and the salt dependence becomes weaker with increased separation of charged residues (Chang et al., 2017; Lin et al., 2016, 2018). Taken together our results suggest that the lack of CTL sequences above and below the thresholds shown in Figure 1C derives from a combination of considerations including maintaining FtsZ concentrations within the cell and minimizing tail-to-tail associations that lead to strong albeit aberrant multivalent interactions amongst segregated blocks of charge within CTTs.

Deletion of the FtsZ CTT leads to a 3-fold increase in GTPase activity. Thus, the CTT plays an auto-inhibitory role by promoting polymerization and lowering the efficiency of GTP hydrolysis. In this way, although GTP binding is required to promote canonical FtsZ polymerization, the hydrolysis of GTP provides the energy required to bend / deform FtsZ polymers to form Z-rings. Preliminary atomistic simulations of the CTT tethered to the FtsZ core show that the CTP motif makes fluctuating patterns of contacts with the T7 activation loop of the FtsZ core (data not shown). We propose that in addition to stabilizing the protofilaments and reducing subunit turnover by lowering the GTPase activity, the CTL may also enable inhibitory interactions between the CTP and the active site of the enzyme. This auto-inhibition could further reduce the GTPase activity and promote polymerization by weakening the dissociation of FtsZ subunits. Similar auto-inhibitory roles have been ascribed to disordered tails that are tethered to ordered domains in a range of different systems (Trudeau et al., 2013).

Recently, Petrovska et al. also showed that metabolic enzymes in yeast form filaments under conditions of stress as a strategy to inactivate these enzymes (Petrovska et al., 2014). Our findings are complimentary to those of Petrovska et al. inasmuch as filament formation weakens GTPase activity of FtsZ (Figure 4G). However, in the bacterial context, the weakening of enzymatic activity ensures robust FtsZ polymerization, which provides the basis for ring formation and serves as a generator of multivalency of CTP motifs that helps coordinate membrane anchoring and a network of interactions with effector proteins and regulatory molecules. The overall picture that emerges is of a multi-functional, multi-purpose CTT with definable features appear to govern the evolution of its amino acid sequence. The sequence complexity has to be sufficiently high vis-à-vis a random control to ensure cellular stability of FtsZ. The CTT has to enable auto-regulatory functions of suppressing GTP hydrolysis by promoting polymerization in the presence of GTP (Figure 7). It is worth noting that polymers of FtsZ will be defined by a multivalency of CTP motifs. From a functional standpoint, this should provide the basis for membrane anchoring and mediating a full complement of protein-protein interactions. As far as cell division and cell growth are concerned, the intra-CTT sequence segregation of oppositely charged residues cannot cross a defined threshold value. If this threshold value is crossed, then the regulatory tail-mediated interactions can be overwhelmed by GTP-independent tail-to-tail interactions. These strong tail-mediated will promote alternative assemblies that compromise canonical FtsZ polymerization and Z-ring formation while conferring a dominant negative phenotype.

**Figure 7:**
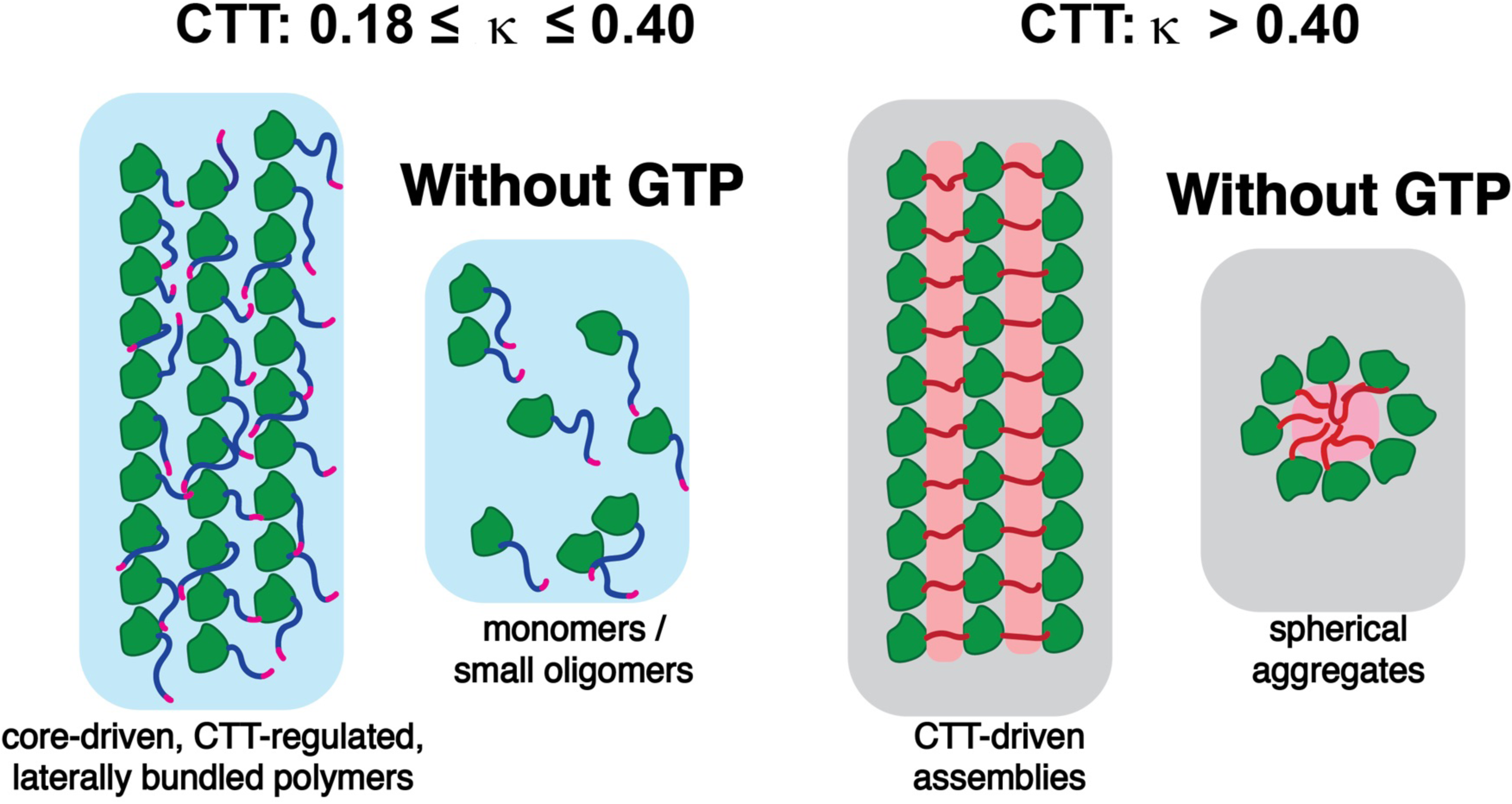
Illustration of the different assemblies driven by GTP versus by the CTT as a function of CTT κ. In the presence of GTP, FtsZ variants with κ values within the evolutionary bounds polymerize and associate laterally via a GTP-dependent and CTT-mediated process. Polymerization and bundling increases the effective valence of the CTP, shown in pink. In the absence of GTP, FtsZ subunits exist primarily as monomers and small oligomers. As CTT κ is increased above 0.4, the interactions amongst CTTs alter the FtsZ assembly mechanism. In the presence of GTP, high κ FtsZ variants form large linear polymers that become laterally zippered through stable CTT-CTT interactions, forming a train track-like morphology thereby inhibiting GTP hydrolysis and slowing down subunit turnover. In the absence of GTP, FtsZ interactions amongst CTTs lead to the formation of spherical assemblies that are stabilized in higher salt concentrations.

A defining feature of canonical FtsZ polymers that form in the presence of GTP and apparently aberrant tail-driven assemblies that form in the absence of GTP is the multivalency of CTT sequences. Whilst canonical FtsZ polymers may be thought of as bottlebrushes with CTTs forming the polymer brushes (Figure 7), the spherical aggregates may be thought of as inverted versions of “hairy spherical colloids” (Lindenblatt et al., 2001) (Figure 7) with the CTTs forming electrostatically stabilized cores and the FtsZ cores making up the corona, although this hypothesis needs to be tested experimentally. Both forms of assemblies are likely candidates for driving phase separation and gelation through the multivalency of CTTs and / or FtsZ cores. We raise this point because multivalency of associative motifs / regions is a defining feature of proteins that drive phase separation and gelation (Banani et al., 2017; Banjade et al., 2015; Harmon et al., 2017; Li et al., 2012). It is worth noting that the bacterial cytoplasm has been shown to form structured glasses (Parry et al., 2014), which would involve physical crosslinking *i*.*e*., gelation of multivalent molecules. We conjecture that the multivalency engendered by polymerization of molecules such as FtsZ with their folded cores and disordered tails might drive the gelation and / or glass-like properties of the bacterial cytoplasm.

Our findings suggest that the CTT / CTL encoded effects cannot be too weak (low κ) or too strong (κ > 0.4); instead, they have to be just right because the CTT plays multiple regulatory roles. In addition, the sequence complexity of the CTT also has to be above a threshold value to ensure protein stability in the cellular setting. We refer to this combination of requirements as a “Goldilocks precept” for the evolution of sequence features within IDPs / IDRs. Similar results have been uncovered from a deep mutational scan of the transactivation domain of Gcn4, which is an essential transcription factor in yeast (Staller et al., 2018). Our investigations open the door for systematic high-throughput experiments guided by recent computational advances (Harmon et al., 2016) that enable the design of CTT sequences with bespoke sequence complexities, lengths, amino acid compositions, and charge patterning. These methods can be applied to other GTPases with N- and C-terminal IDRs of GTPases in order to uncover the robustness of the Goldilocks precept that appears to underlie the evolutionary selection of IDPs / IDRs.

### Experimental Procedures

#### General methods

*B*. *subtilis* strains expressing CTL variants of FtsZ were derived from PAL 2084 and were grown in 0.5% xylose to induce wild-type expression or 0.1 mM IPTG to induce CTL variant expression. Vent DNA polymerase was used for PCR (New England Biolabs). All restriction enzymes were ordered from New England Biolabs. All genomic DNA extractions were performed using the Wizard Genomic DNA Purification Kit (Promega). Plasmid preparations were made using the NucleoSpin Plasmid Kit (Macherey-Negel). Gel/PCR purifications were performed using the NucleoSpin Gel and PCR Clean-up Kit (Macherey-Negel). T4 DNA ligase was used for ligations (New England Biolabs).

#### Cloning CTL variants

The CTL variant strains were constructed as described previously (Buske and Levin, 2013). The bacterial strains and plasmids used in this study are listed in **Table S1** of the supplemental material. Synthetic double-stranded oligonucleotides of the CTL variants were ordered from Integrated DNA Technologies, digested using restriction enzymes, and ligated into pPJ19, which contains FtsZ, under the control of the Pspac promoter that is inducible with 0.1 mM IPTG, with restriction sites flanking the CTL. A BamHI site after amino acid 315 and a XmaI site before residue 366 result in the insertion of amino acid pairs GS and PG N- and C-terminal to the CTL, respectively. The plasmid was transformed into PAL 644, a strain of *E*. *coli* derived from PAL930. PAL 644 contains the low copy plasmid pBS58 expressing *E*. *coli* ftsQAZ, which allows the sub-cloning of *B*. *subtilis* FtsZ. FtsZ was amplified in pPJ19, the product of which was restriction digested, purified, and ligated into pDR67. The multiple cloning site in the vector pDR67 contains the 5’- and 3’-ends of the amyE gene on either side, which allows the insertion of FtsZ into the amyE locus by homologous recombination. The purified plasmid was transformed into PAL 522, a derivative of the JH642 wild-type strain of *B*. *subtilis*. Genomic DNA was purified and transformed into MEO 1, a derivative of PAL 2084 containing a copy of wild-type FtsZ under the control of the Pxyl promoter, inducible with 0.5% xylose. The cells were made competent and transformed with purified genomic DNA from PAL 2084, which knocks out the chromosomal WT copy of FtsZ. Plasmids were verified by restriction digests and sequencing.

#### Immunoblotting

Immunoblotting was performed as described previously (Weart and Levin, 2003). Cells were grown overnight in Luria-Bertani (LB) medium at 37°C with 100 μg/mL ampicillin, 100 μg/mL spectinomycin, 5 μg/mL chloroamphenicol and 0.5% xylose. They were then back-diluted 1:100 and grown in 0.5% xylose until the cells reached mid-log phase. The cells were then washed twice with LB, diluted 1:100, and grown to mid-log phase in 0.1 mM IPTG. The cells were lysed with lysozyme and detergent. Loading was normalized to the OD_600_ at sampling. The blot was probed using affinity-purified polyclonal rabbit anti-FtsZ antibodies and goat anti-rabbit antibodies conjugated to horseradish peroxidase (Jackson ImmunoResearch Laboratories). Immunoblots were developed using the ECL Western Blotting detection reagents (GE Healthcare) and visualized with the luminescent image analyzer ImageQuant LAS 4000 mini (GE Healthcare).

#### Growth Curves

Cells were grown overnight under the same media conditions as the immunoblots in 0.5% xylose, back-diluted 1:100 and grown in 0.5% xylose until the cells reached mid-log phase. The cells were then washed twice with LB, diluted to OD_600_ 0.004, and grown in 0.1 mM IPTG. Starting 1 hour after induction, the OD600 was measured every 30 minutes for 4 hours.

#### Immunofluorescence microscopy

Immunofluorescence microscopy was performed as described previously (Buske and Levin, 2012). Cells were grown using same media conditions overnight in 0.5% xylose, back-diluted 1:100 and grown in 0.5% xylose until the cells reached mid-log phase. The cells were then washed twice with LB, diluted 1:100, and grown in 0.1 mM IPTG for 5 generations (∼2.5 hours). The cells were harvested and fixed with 16% paraformaldehyde/0.5% gluteraldehyde. The cells were lysed with 2 mg/mL lysozyme. FtsZ was detected with affinity-purified polyclonal rabbit anti-FtsZ serum in combination with goat antirabbit serum conjugated to Alexa488 (Life Technologies). Cell walls were stained with wheatgerm agglutinin conjugated to tetramethylrhodamine, and DNA was stained with DAPI. Slides were visualized with an Olympus BX51 microscope with Chroma filters and a Hamamatsu OrcaERG camera and processed using Openlab version 5.2.2 (Improvision) and Adobe Photoshop CS version 8.0 (Adobe Systems). The cell length/Z-ring (L/R) ratio was calculated as described by Weart et al. 2007. The L/R ratio was calculated as the sum total cell length of a population of cells divided by the total number of Z-rings in that population.

#### Atomistic Simulations of CTT Sequence Variants

All-atom Monte Carlo simulations were performed using the ABSINTH implicit solvent model and forcefield paradigm as made available in the CAMPARI simulation package (http://campari.sourceforge.net) (Radhakrishnan et al., 2012; Vitalis and Pappu, 2009a; Vitalis and Pappu, 2009b). Simulations utilized the abs_3.2_opls.prm parameter set in conjunction with optimized parameters for neutralizing and excess Na^+^ and Cl^−^ ions (Mao and Pappu, 2012). Simulations were performed using a spherical droplet with a diameter of 285 Å with explicit ions to mimic a concentration of 10 mM NaCl. Temperature replica exchange Monte Carlo (T-REMC) (Sugita and Okamoto, 1999) was utilized to improve conformational sampling. The temperature schedule ranged from 280 K to 400 K. Ensembles corresponding to a temperature of 310 K were used in the analysis reported in this work. Three independent sets of T-REX simulations were performed for each CTT sequence. In all, the ensembles for each CTT sequence were extracted from simulations, where each simulation deploys 4.6 × 10^7^ Monte Carlo steps. In each simulation, the first 10^6^ steps were discarded as equilibration. Simulation results were analyzed using the MDTraj (McGibbon et al., 2015) and CTraj routines that are available at http://pappulab.wustl.edu/CTraj.html.

#### Protein Purification

FtsZ variants were cloned into the pET-21b(+) expression vector through *E. coli* strain AG1111. The resulting plasmids were mini-prepped and freshly transformed into C41(DE3) cells and made into glycerol stocks. 500 mL of LB medium was inoculated 1:100 with an overnight culture. Cells were grown at 37°C until A_600_ ∼0.6-0.8, and then cells were induced with isopropyl IPTG to a final concentration of 1 mM. Cells were grown for an additional 4h at 37°C, and then the cells were harvested by centrifugation, and the cell pellets were stored at −80°C. Purification was carried out as previously described (Buske and Levin, 2012). Peak fractions were analyzed by SDS-PAGE and pooled together, and dialyzed overnight in 1 L of FtsZ dialysis buffer (50 mM MES 50 mM KCl, 2.5 mM MgCl_2_, 1 mM EGTA, 10% sucrose, pH 6.5) Protein preparations were concentrated, separated into aliquots, and flash frozen on liquid N_2_, and stored at –80°C. Prior to use, FtsZ aliquots were thawed on ice, well mixed, and the concentration was quantified using Pierce 660nm assay with tubulin as a standard (Thermo Fisher Scientific).

#### 90° Light Scattering Assay

Assembly reactions contained 5 μM FtsZ in an MES buffer solution (50 mM MES, 2.5 mM MgCl2, 1 mM EGTA, 10% sucrose, pH 6.5, with salt concentrations varying from 50-200 mM KCl as specified). Measurements were recorded every quarter of a second at 30°C. A 1-minute baseline was established prior to adding 1 mM GTP to the reaction. A minimum of three trials was conducted per salt concentration for each variant. All data were collected and exported into Microsoft Excel, and the subsequent analysis was performed using Matlab. The average baseline was subtracted from each data point.

#### Transmission Electron Microscopy (TEM)

Samples were prepared in conditions mimicking the light scattering assays, with a lower concentration of FtsZ (2.5 μM). Prior to preparing the copper grids, each sample was incubated for 10 minutes in the presence of 1 mM GTP to allow for adequate assembly. Each sample was stained 3x with 2% uranyl acetate for 20-seconds each, with the solution wicked away and followed by a 10 second wait time in between stains. Samples were visualized using a FEI Transmission Electron Microscope.

#### GTPase Assay

GTP hydrolysis activity was monitored using a coupled GTPase assay (Ingerman and Nunnari, 2005). Using a 96-well plate reader, the assay was conducted in the same buffer conditions as the light scattering assay (5 μM FtsZ WT / variant, and 1 mM GTP), and included 1 mM phosphoenolpyruvate, 250 μM NADH, and 40 units/ml of both lactose dehydrogenase and pyruvate kinase. From the following equation, the linear decline of NADH absorbance at 340 nm was monitored over 30 minutes. The steepest decline rate for a 5 minute consecutive stretch was related to the GTPase activity by the following manipulation of Beer’s law, which yields:
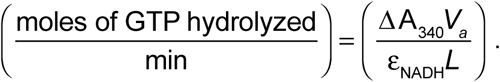
Here, ΔA_340_ is the slope of the decline, ε_NADH_ is the extinction coefficient for NADH at 340 nm (6220 M^−1^cm^−1^), *L* is the path length of the cuvette (0.401 cm), and *V_a_* is the observation volume (150 μL). Each trial was performed in triplicate.

#### Image Processing

FtsZ TEM images were analyzed using ImageJ software. Length scale comparisons were made under the assumption that each amino acid in the linker is approximately 0.38 nm in length, as generalized for an extended polypeptide.

## Acknowledgments

The US National Science Foundation supported this work through grants MCB-1614766 and MCB-0718924 to RVP. We are grateful to our current and former colleagues Jeong-Mo Choi, Rahul Das, Martin Fossat, Tyler Harmon, Jared Lalmansingh, and Kiersten Ruff for helpful discussions.

## Supplemental Material

**Figure S1:**
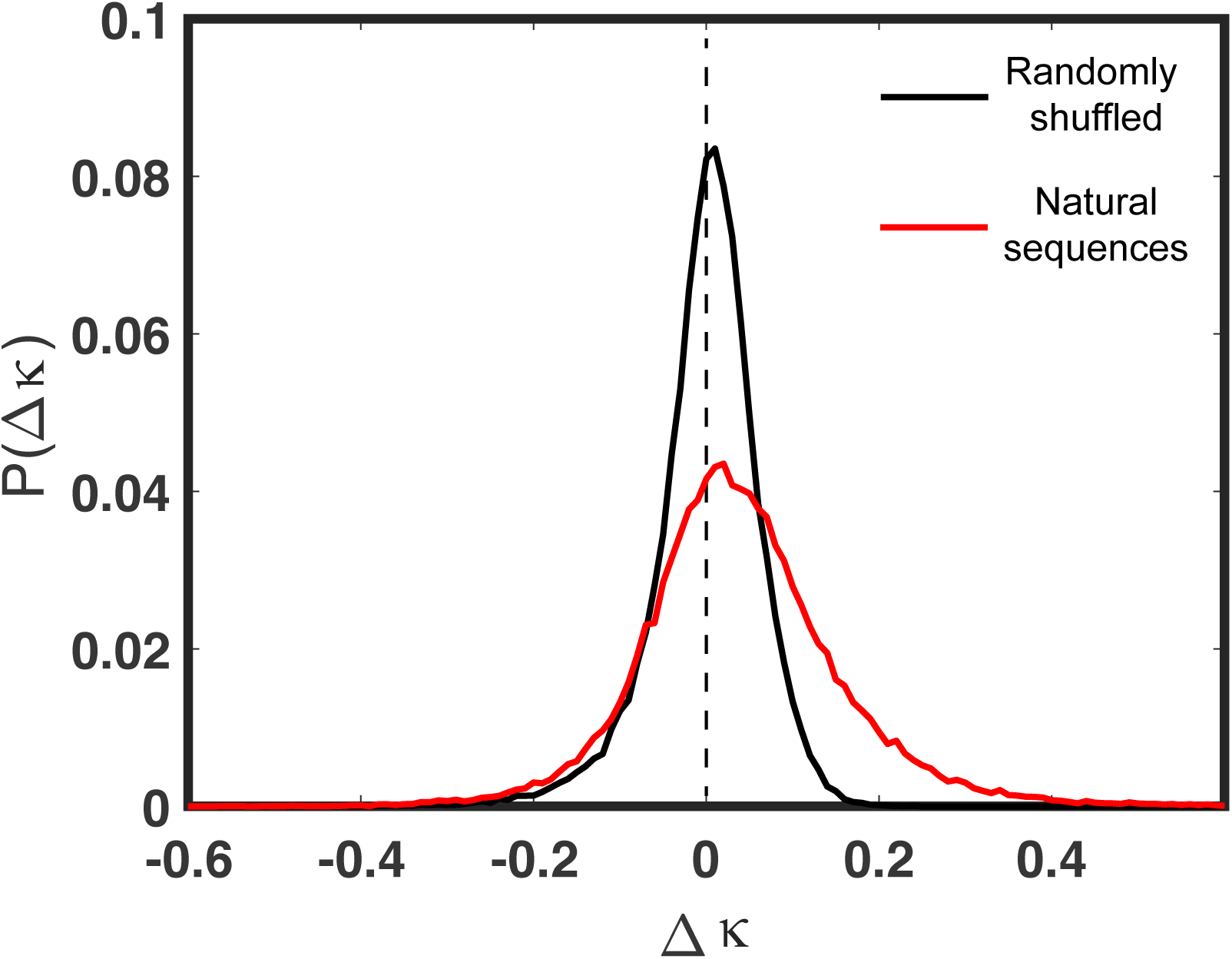
Naturally occurring sequences with an FCR ≈ 0.3 have a more positive κ value than would be expected by random chance. We identified all predicted IDRs from the human protein within the bounds 0.27 < FCR < 0.33 that were thirty residues or more in length (4755 IDRs). For each IDR, we calculated the expected κ (〈κ〉) value associated with a sequence of that composition in the absence of any selective pressure performing multiple random shuffles and assessing the mean value. Finally, for each naturally occurring IDR (WT), we calculated the Δκ, where Δκ = κ^WT^ – 〈κ〉. If Δκ > 0 then the naturally occurring sequence has a higher κ than expected, while if Δκ < 0 the naturally occurring sequence has a lower κ than would be expected, and the distribution of Δκ was plotted as the red curve. The same analysis was performed for randomly shuffled sequences instead of naturally occurring sequences to assess the intrinsic variance in the data, as shown in the black curve. Many sequences showed a bias for higher than expected κ values, although a small number of sequences showed a bias for lower than expected κ values.

**Figure S2:**
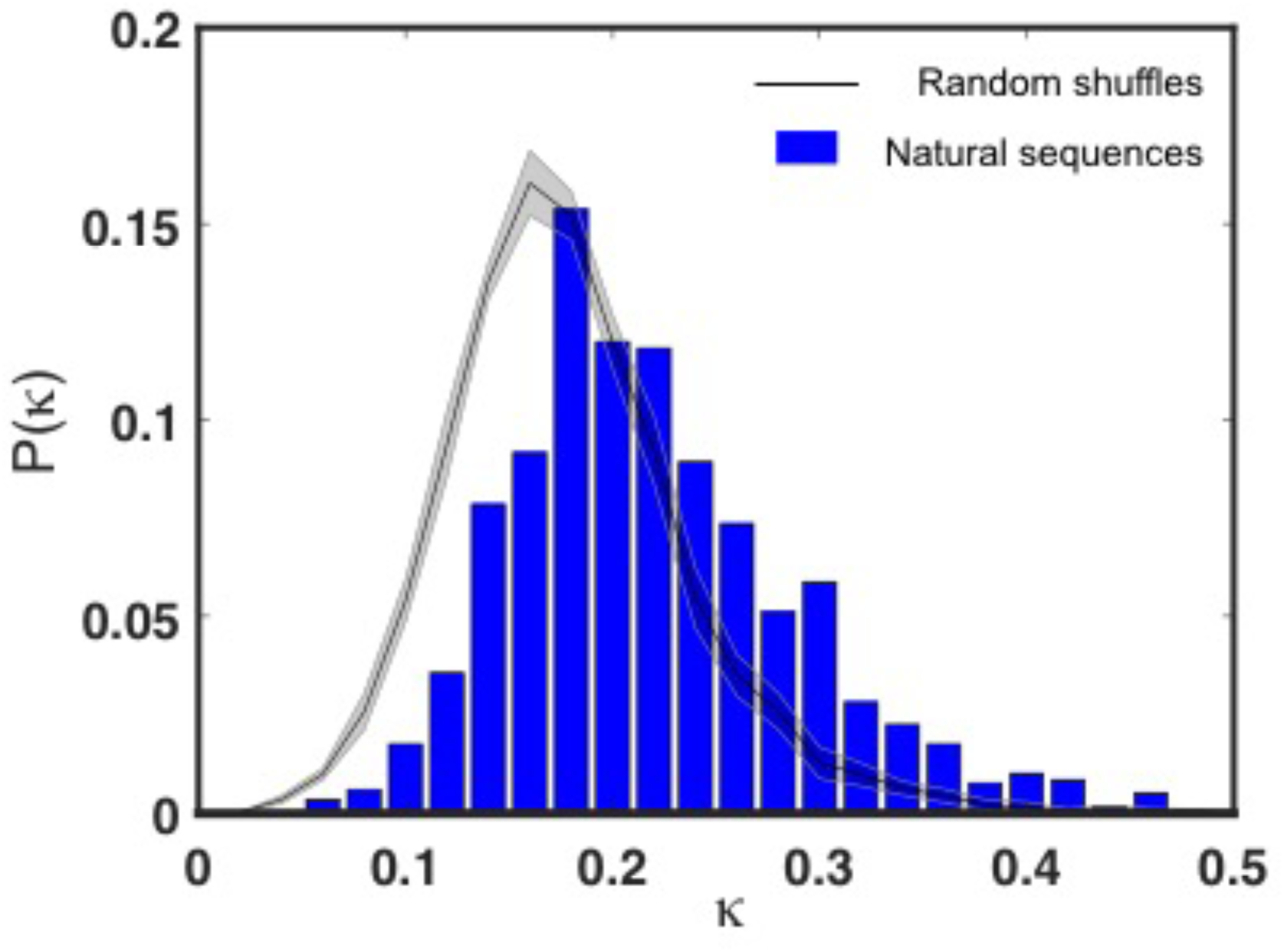
Naturally occurring FtsZ CTT sequences are biased towards larger κ than expected by random chance. We calculated the κ value associated with 1209 orthologous FtsZ CTT sequences. The associated distribution of κ values is shown in blue. For each sequence, we then shuffled the sequence multiple times to generate a set of randomly permuted CTT sequences. The average distribution (and associated variance) is shown in the black and grey curves. These data demonstrate that naturally occurring FtsZ CTT sequences are biased to more positive κ values than expected by random chance.

**Figure S3:**
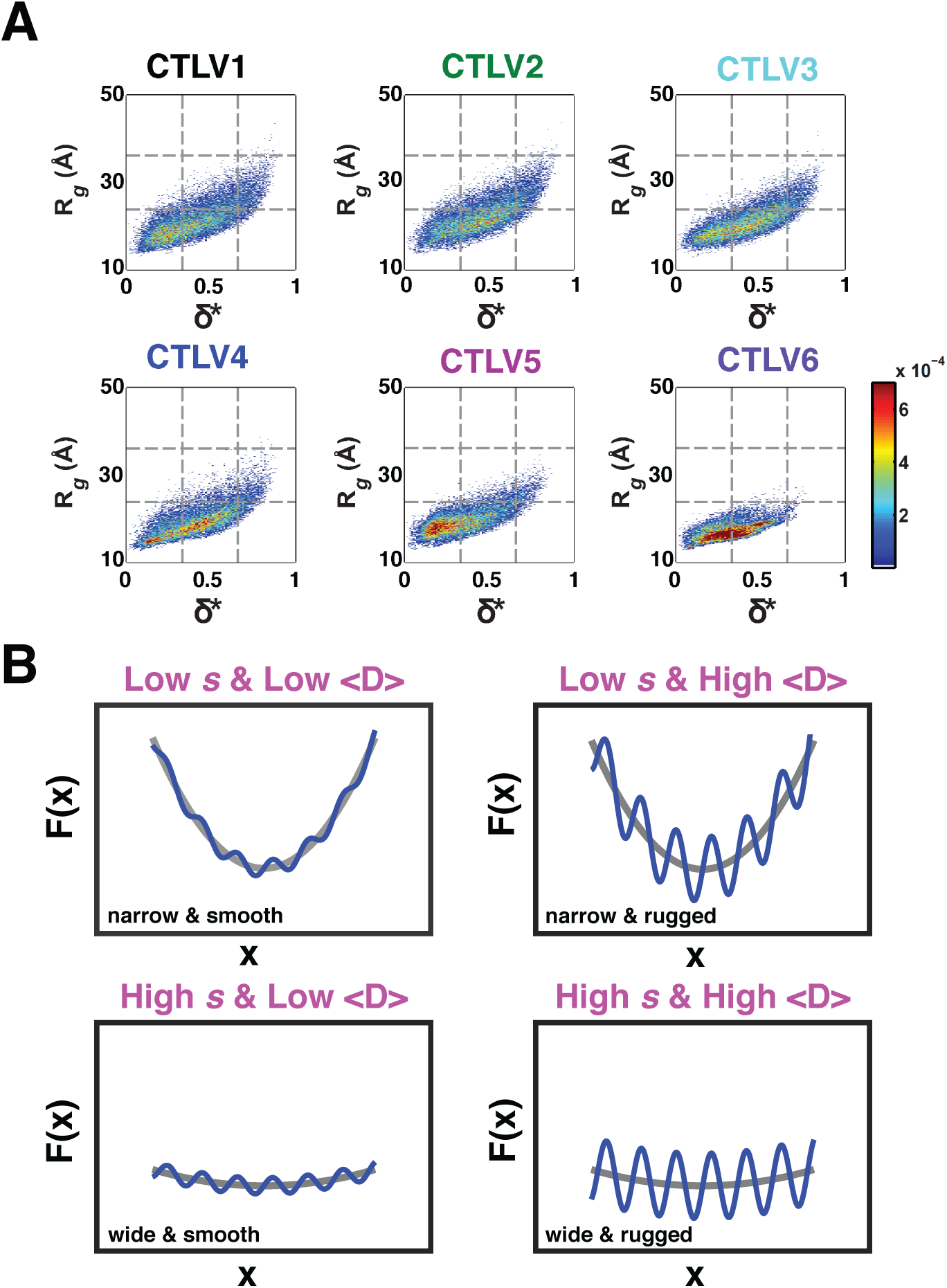
**(A)** Conformational distributions from atomistic simulations of CTT sequences for all six CTT variants. Results obtained with the same conditions as Figure 2D. (B) Conformational landscape topology and breadth are outlined by contributions from sizes, shapes, and backbone dihedral angles. Landscapes can be narrow and smooth (low *s* and *D*), narrow and rugged (low *s* and large *D*), wide and smooth (high *s* and low *D*), or wide and rugged (high *s* and *D*).

**Figure S4:**
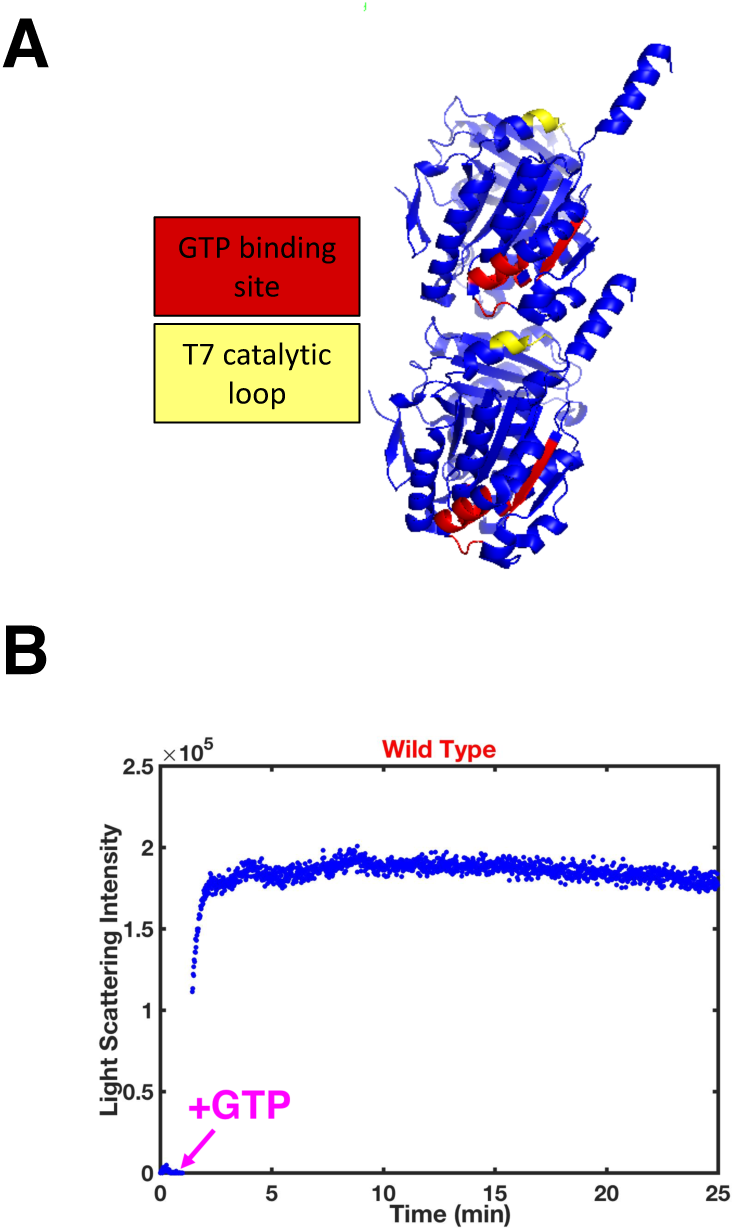
FtsZ forms protofilaments and polymerization is sensitive to the presence of GTP. (A) FtsZ is a GTPase whose active site is formed by the interface between two FtsZ monomers. The cartoon representation, generated using the VMD software package, is based on the coordinates for FtsZ dimers obtained from the protein data bank (PDB id: 2vxy). In each subunit, residues that make up the GTP binding site are colored in red whereas residues that make up the T7 activation loop are colored in yellow. **(B)** Data from a typical light scattering GTPase polymerization assay, shown here for the WT FtsZ. The arrow marks the time point for introduction of GTP into the solution mixture. The blue trace represents data points collected as a function of time. The intensity of scattered light is sensitive to the sizes of aggregates formed in solution. The larger the aggregates, the greater the intensity of scattered light and aggregates can be a combination of protofilaments and bundled polymers.

**Figure S5:**
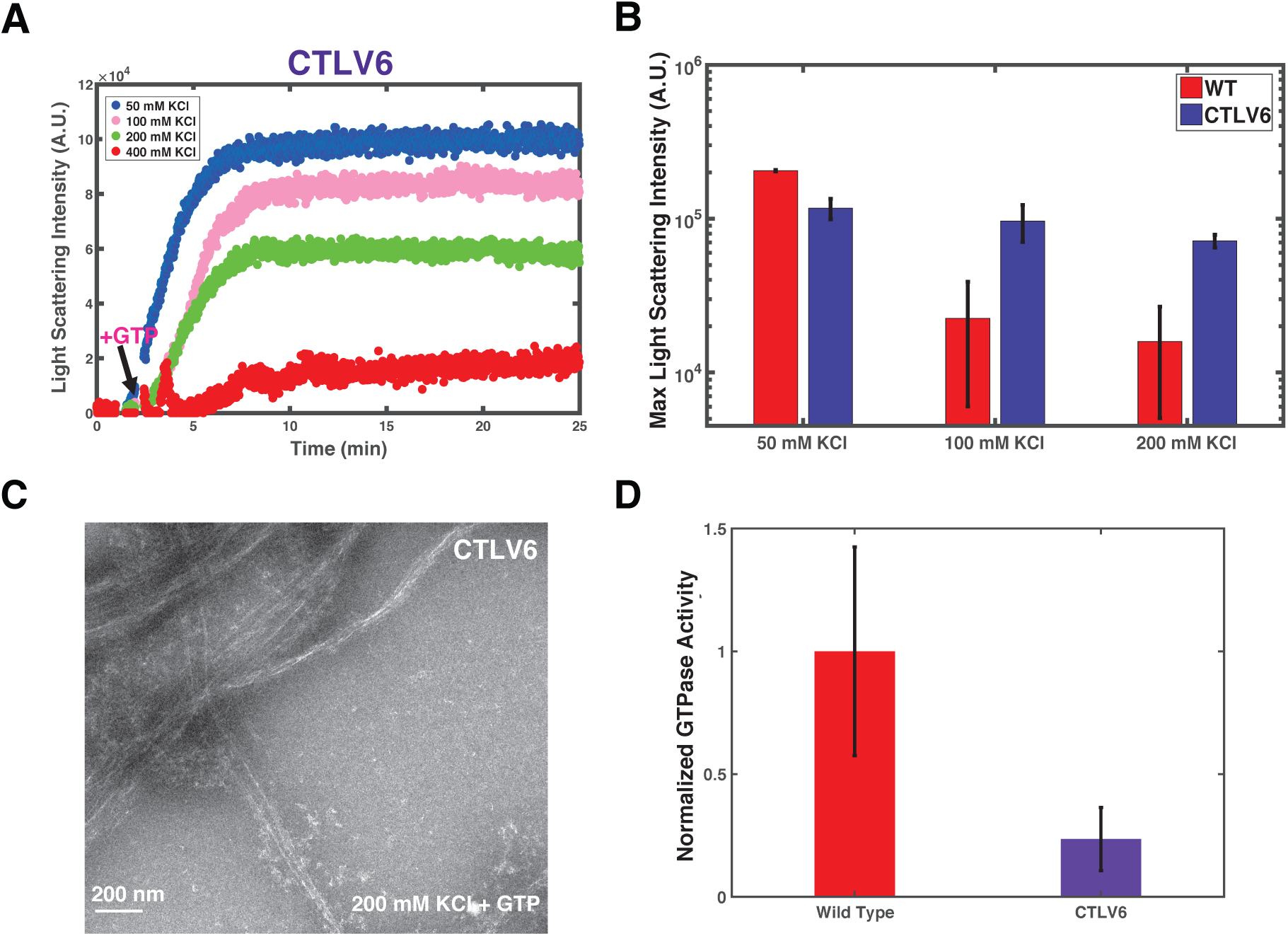
(A) CTLV6 assembly is incredibly robust in the presence of increasing salt concentration. (B) The relative maximum scattering intensities for 50, 100, and 200 mM salt concentrations illustrate the robustness of CTLV6 assembly despite the increase in salt concentration. (C) EM images show that at 200 mM KCl the CTLV6 morphology is maintained. (D) CTLV6 shows minimal GTP hydrolysis activity when compared to the WT.

**Figure S6:**
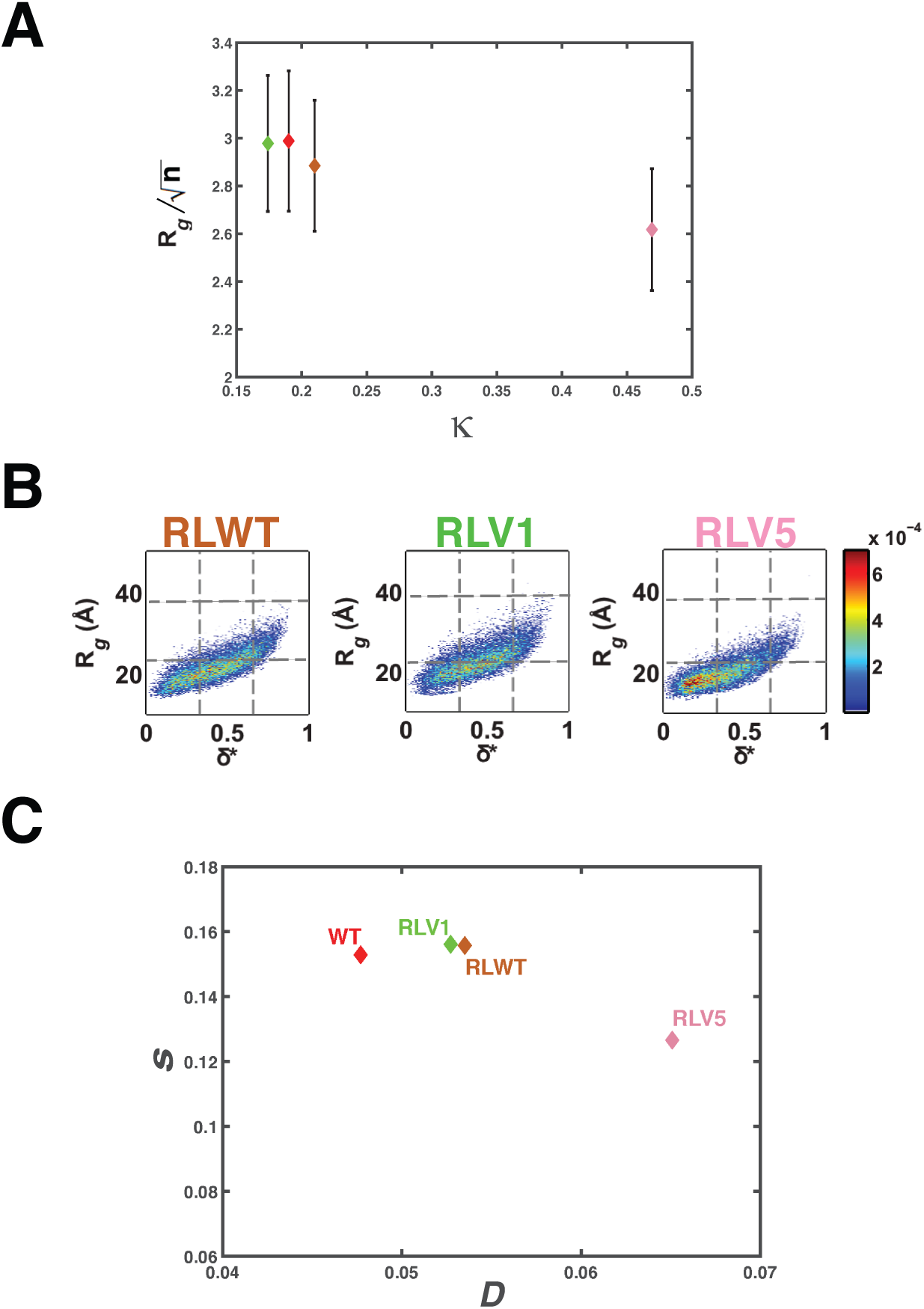
(A) Plots of length normalized values of the mean radii of gyration for each of the reduced library CTT variants. These values were obtained as in Figure 2C. (B) Conformational distributions from atomistic simulations of CTT sequences for the reduced library sequences in terms of size (R_g_) and shape (asphericity). (C) Plot of the values of D and *s* for the WT CTT and each of the designed reduced library CTT variants.

